# Isoform-specific targeting properties of the protocadherin CDHR5 control its apical delivery to promote brush border assembly

**DOI:** 10.1101/2023.02.22.529570

**Authors:** Samaneh Matoo, Maura J. Graves, Myoung Soo Choi, Rawnag A. El Sheikh Idris, Prashun Acharya, Garima Thapa, Tram Nguyen, Sarah Y. Atallah, Ashna K. Tipirneni, Phillip J. Stevenson, Scott W. Crawley

## Abstract

Transporting epithelial cells of the gut and kidney interact with their luminal environment through a densely-packed collection of apical microvilli known as the brush border. Proper brush border assembly depends on the intermicrovillar adhesion complex (IMAC), a protocadherin-based adhesion complex found at the distal tips of microvilli that mediates adhesion between neighboring protrusions to promote their organized packing. Loss of the IMAC adhesion molecule Cadherin-related family member 5 (CDHR5) correlates with poor prognosis of colon cancer patients, though the functional properties of this protocadherin have not been thoroughly explored in relevant cell systems. Here, we show that the two dominant CDHR5 splice isoforms expressed in enterocytes interact to form an apparent *cis*-oligomer that is competent to target to the apical domain to drive microvillar elongation. The two isoforms exhibited distinct sequence-dependent apical targeting properties, with one isoform requiring its cytoplasmic tail. Library screening identified the Ezrin-associated scaffolds EBP50 and E3KARP as cytoplasmic binding partners for CDHR5. Consistent with this, loss of EBP50 disrupted proper brush border assembly with cells exhibiting markedly reduced apical IMAC levels. Together, our results shed light on the apical targeting determinants of CDHR5 and further define the interactome of the IMAC involved in brush border assembly.

## INTRODUCTION

During their terminal differentiation, transporting epithelial cells of the gut and kidney assemble a dense array of actin-based microvilli on their apical surface that are specialized to mediate solute transport (1). Collectively, these microvilli are referred to as a brush border (BB) and act to amplify the apical surface area of the cell that is in contact with the external environment. As part of this optimization, these epithelial cells organize their microvilli into hexagonal arrays: a geometric pattern that allows for the maximum number of microvilli to occupy the apical surface of each cell to ensure peak solute transport. Indeed, each transporting epithelial cell assembles an organized collection of ∼1000 apical microvilli in their mature state. Perturbations to BB structure can be life threatening unless treated, as seen with infections of the gut by the attaching/effacing microbe enterohemorrhagic E. coli (2) and Antibrush Border Antibody Disease in which autoantibodies damage the BB of the kidney (3).

Previous studies focused on understanding the underlying mechanism of BB development identified the intermicrovillar adhesion complex (IMAC), a protocadherin-based complex found at the distal tips of BB microvilli that creates physical linkages that connect neighboring microvilli together to control their organization (Fig. S1A) (4). These ‘intermicrovillar adhesion links’ are a *trans*-heterophilic adhesion complex between Cadherin-related family member 2 (CDHR2; also known as protocadherin-24) and Cadherin-related family member 5 (CDHR5; also known as mucin-like protocadherin) (Fig. S1A)(4). By forming physical connections between the distal tips of neighboring microvilli, the IMAC organizes these microvilli into discrete ‘tee pee-like’ apical clusters during the early stages of BB assembly. As these clusters grow larger with the addition of new microvilli, they eventually amalgamate to form a unified, highly organized BB. Both *in vitro* and *in vivo* studies have shown that IMAC-mediated intermicrovillar adhesion plays a critical role in proper BB assembly (4-7). CDHR2 knockout (KO) mice exhibit malformed enterocyte BBs, having short microvilli that do not attain their normal ordered dense packing (8). As a consequence of this, CDHR2 KO mice have a lower body weight compared to heterozygous littermates, likely due to a reduced functional capacity of their intestinal tissue in nutrient absorption. Studies revealed that proper targeting and function of CDHR2 requires association with a cytoplasmic complex comprised of two scaffolding molecules, Ankyrin repeat and sterile α-motif domain containing 4B (ANKS4B) and USH1C (also known as harmonin), a myosin motor protein Myosin-7b (Myo7b), and the myosin light chain Calmodulin-like protein-4 (CALML4) (Fig. S1A)(4,6,7). In contrast, little is currently known about the functional properties of CDHR5, as well as its cytoplasmic binding partners that may contribute to its cellular role.

CDHR5 was originally discovered in a screen for proteins involved in branching morphogenesis that occurs during development of the kidney (9,10). Subsequently, CDHR5 was also identified to be highly expressed in the gut as two distinct splice isoforms (10,11) and was found to be lost in most colorectal cancer tissue samples analyzed for the protein (12). Epigenetic silencing through methylation of the promoter region of CDHR5 is thought to be one mechanism that mediates downregulation of this gene during colorectal cancer development (13). However, how CDHR5 helps maintain the normal integrity of the intestinal epithelium and guards against tumor formation is currently unclear. In this study, we explored the functional properties of CDHR5 using cultured proximal tubule kidney epithelial cells and enterocytes as cell models that generate an apical BB. Using these systems, we questioned whether the two dominant splice isoforms of CDHR5 exhibited different targeting properties, and if so, what sequence features of these isoforms are important for these distinct properties. These studies led us to further screen for potential cytoplasmic binding partners for CDHR5 to understand how they may contribute to its function. Together, our results shed light on how CDHR5 is delivered to and promotes formation of an apical BB, a critical interface found between transporting epithelia of the gut and kidney and their luminal environments.

## RESULTS

### The splice isoforms of CDHR5 exhibit different targeting properties

To begin to examine the properties of CDHR5 in transporting epithelia, we investigated its localization in duodenal and colon tissue samples from mice. We used the actin crosslinking protein Villin as a counterstain to visualize the BB in tissue samples. CDHR5 exhibited prominent targeting to the distal tips of duodenal BB microvilli, consistent with it being a component of the IMAC (Fig. 1A; arrowheads). Within colon tissue sections, CDHR5 signal appeared more uniformly distributed across the microvillar axis, though areas of enrichment towards the distal tips of these shorter microvilli could also be seen (Fig. 1B; arrowheads). Interestingly, distinct subapical CDHR5 puncta were often seen just below the BB for both tissue samples (Fig. 1A,B; arrows). Similar subapical puncta were not previously observed for other known IMAC components (Fig S1B,C; CALML4 provided as example), suggesting that they may be a specific property of CDHR5. We turned to examine the localization of CDHR5 in CACO-2_BBE_ cells, a human enterocyte model system that utilizes the IMAC to construct a near tissue-like apical BB when allowed to polarize for an extended period of time in cell culture (4,14). Staining for CDHR5 in these polarized cells revealed areas of the monolayer that had signal that appeared ‘trapped’ internally below the apical BB (Fig. 1C; arrows). We compared this to polarized CACO-2_BBE_ monolayers stained for CDHR2, the adhesion binding partner for CDHR5 within the IMAC (4). In contrast to CDHR5, CDHR2 did not display a similar population of internally trapped protein (Fig. S1D). Measuring the BB to cytoplasmic ratio of signal for both IMAC protocadherins revealed that CDHR2 exhibited a significantly higher efficiency of BB targeting compared to CDHR5 (Fig. 1D). To guard against the potential of off-target labeling, we confirmed the specificity of our CDHR5 antibody by disrupting the CDHR5 transcript in CACO-2_BBE_ cells using a previously validated shRNA (4). Immunoblot for CDHR5 from cell lysates derived from polarized CACO-2_BBE_ monolayers expressing a scramble control detected distinct bands at ∼70 kDa, ∼100 kDa and ∼200 kDa, while these products were not observed in the CDHR5-targeted shRNA stable cell line (Fig. 2A). In agreement with this, immunostaining using this antibody also failed to detect prominent signal in our CDHR5 KD cell line (Fig S2A).

**Figure 1.**
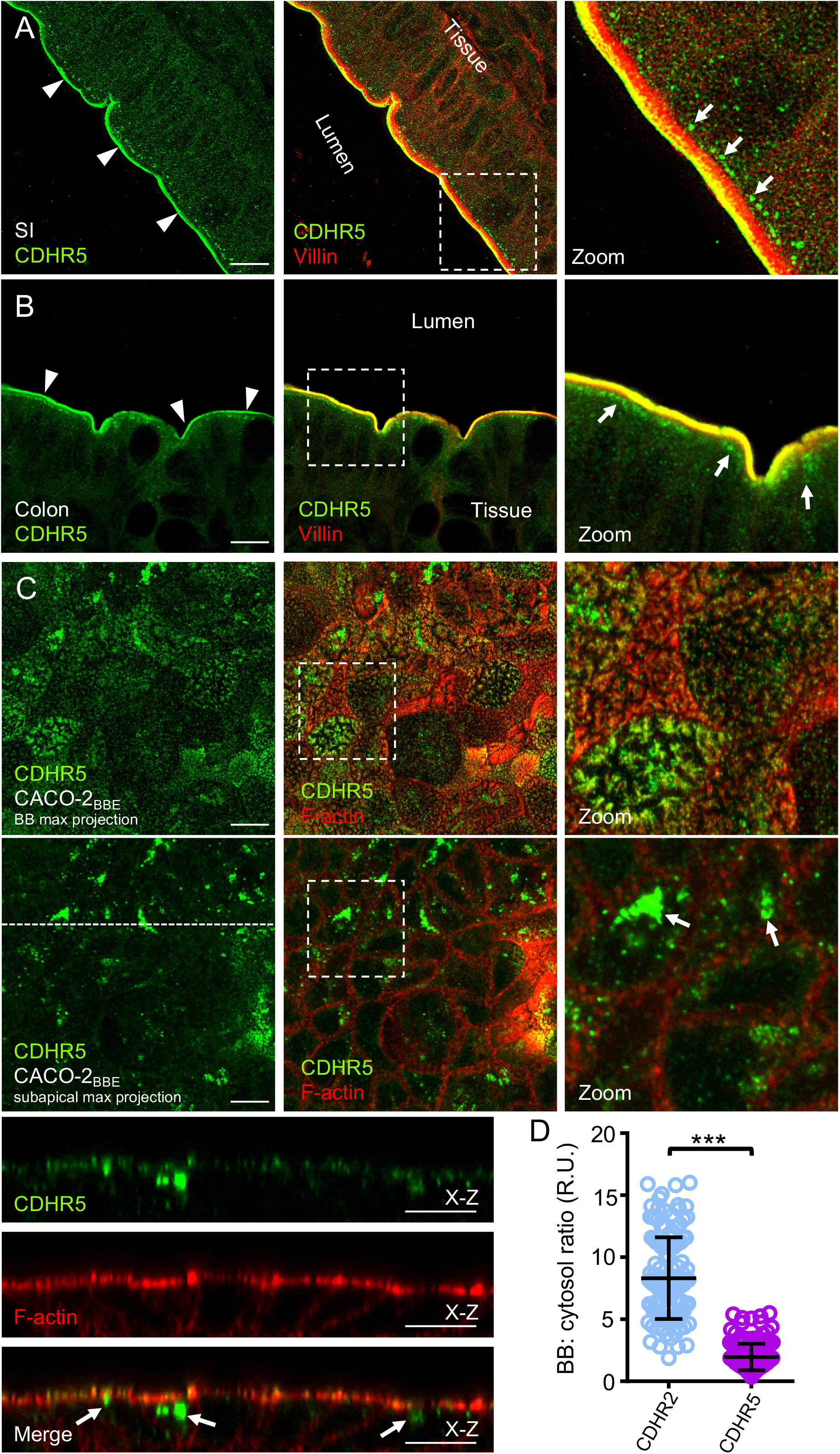
Targeting properties of CDHR5 in intestinal tissue and CACO-2_BBE_ cells. (A+B) Confocal images of mouse small intestine (SI) and colon tissue stained for CDHR5 (green) and Villin (red). Arrowheads denotes localization to the tips of BB microvilli, while arrows point to signal accumulation found in subapical puncta. Boxed regions denote area in zoomed image panels. Scale bars, 10 μm. Unlike other IMAC components, distinct signal of CDHR5 is detected as puncta below the BB. (C) Confocal images of 12-day polarized CACO-2_BBE_ cells stained for endogenous CDHR5 (green) and F-actin (red). Upper panel series show a maximum projection through the full height of the BB, while lower panels show a maximum projection through the same monolayer excluding the apical BB. Arrows shown in the zoom image for the lower panel series point to large subapical and cytoplasmic puncta. Boxed regions denote area in zoomed image panels. Dashed line in the subapical maximum projection indicate the position where the x-z section was taken; Individual channels for the *x-z* section are shown below the *en face* image. Scale bars, 10 μm. Similar to tissue, CDHR5 signal is detected below the BB. (D) Scatterplot quantification of the BB:cytosol ratios of endogenous CDHR5 and CDHR2 signal for 12-day polarized CACO-2_BBE_ cells. Bars indicate mean and SD. R.U = relative units. ***p < 0.0001, two-tailed t test. CDHR2 exhibits a significantly higher efficiency of BB enrichment compared to CDHR5.

**Figure 2.**
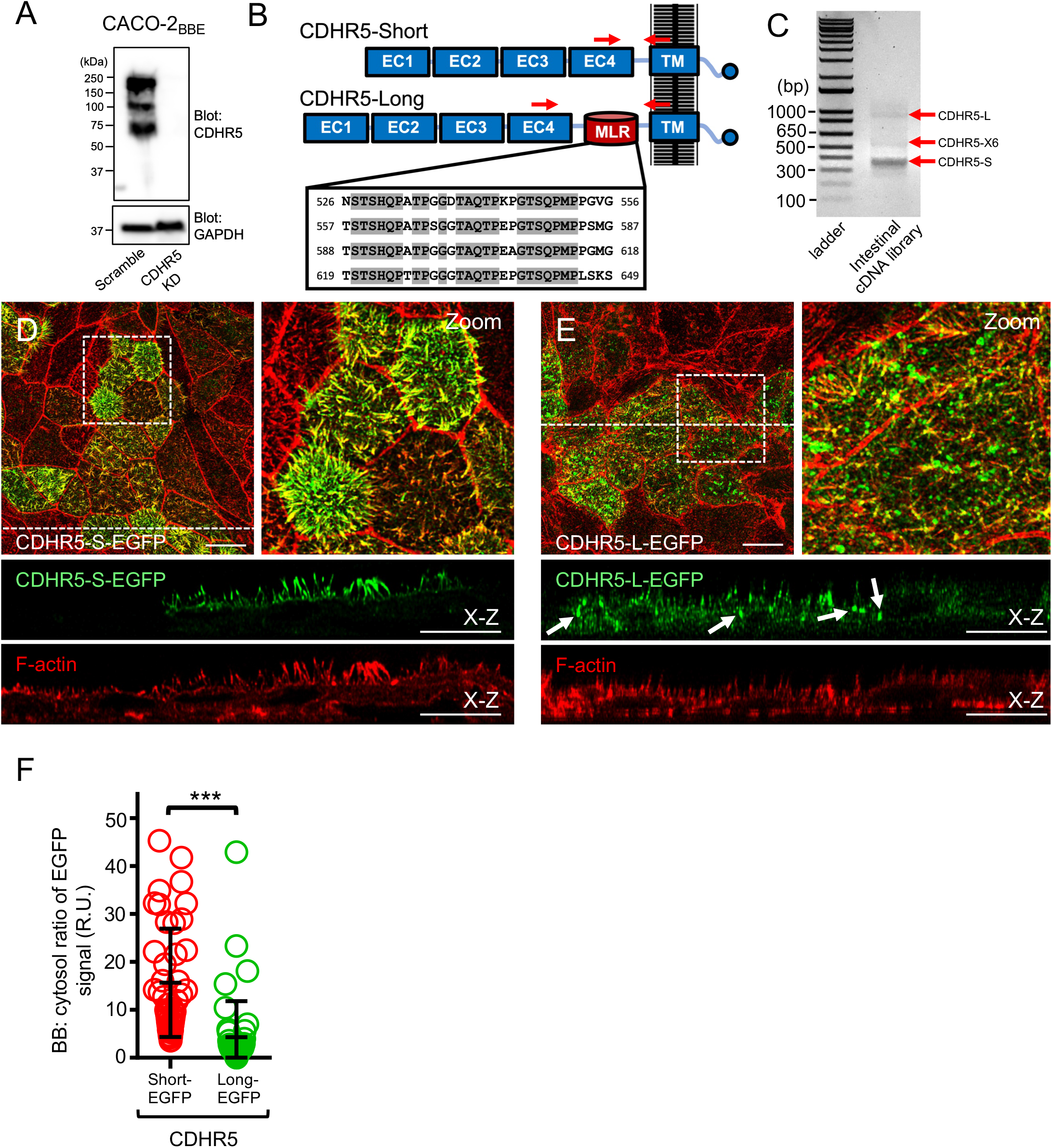
The splice isoforms of CDHR5 exhibit different targeting properties in LLC-PK1-CL4 kidney epithelial cells. (A) Immunoblot analysis of endogenous CDHR5 in lysates from 21-day polarized scramble shRNA control and an shRNA CDHR5 KD stable CACO-2_BBE_ cell line. GAPDH served as a loading control. CDHR5 is detected as three distinct bands in CACO-2_BBE_ cell lysates. (B) Cartoon domain diagram of the two splice isoforms of human CDHR5. Inset panel shows the sequence of the 4X mucin-like tandem repeat found in the MLR domain of the long isoform of human CDHR5. EC=extracellular cadherin repeat; MLR=mucin-like repeat domain; TM=transmembrane domain. Red arrows indicate where the primer set used in (C) anneals to the CDHR5 transcript. (C) A primer set designed to flank the MLR domain of CDHR5 was used for PCR analysis of an intestinal cDNA library. Cloning and sequencing of the products revealed that the band at ∼950 bp corresponds to CDHR5-L (expected size = 946 bp) and the band at ∼350 bp corresponds to CDHR5-S (expected size = 946 bp). The band just above ∼550 bp was identified as a potential new splice isoform of CDHR5 that we designated as CDHR5-X6. (D+E) Confocal images of 4-day polarized LLC-PK1-CL4 cells stably expressing either EGFP-tagged CDHR5-S or CDHR5-L (green) and stained for F-actin (red). Boxed regions denote area in zoomed image panels. Dashed lines indicate the position where the *x-z* sections were taken; Individual channels for the *x-z* sections are shown below the *en face* images. Arrows in the CDHR5-L-EGFP green channel *x-z* section point to subapical and cytoplasmic puncta of signal. Scale bars, 10 μm. CDHR5-L exhibits a lower efficiency of targeting to the BB of LLC-PK1-CL4 cells and can be often found in subapical and cytoplasmic puncta. (F) Scatterplot quantification of the BB:cytosol ratios of EGFP-tagged CDHR5-S and CDHR5-L signal for 4-day polarized LLC-PK1-CL4 stable cell lines. Bars indicate mean and SD. R.U = relative units. ***p < 0.0001, two-tailed t test.

CDHR5 has previously been reported to be expressed in transporting epithelial cells predominantly as two splice isoforms, a short isoform (CDHR5-S; theoretical molecular weight 69.6 kDa) and a long isoform (CDHR5-L; theoretical molecular weight 88.2 kDa) (Fig. 2B)(10,15). The two CDHR5 isoforms are identical in sequence with the exception that the long isoform contains an additional membrane proximal mucin-like repeat (MLR) domain in its extracellular region (Fig. 2B). In humans, the CDHR5 MLR domain is comprised of a 4X tandem repeat sequence of 31 amino acids that is rich in threonine, serine and proline (TSP) residues, similar to a conventional TSP-rich region found in the secreted and transmembrane mucins of the gut (Fig. 2B). We designed primers that flanked the MLR domain and confirmed that transcript for both the CDHR5-L and CDHR5-S isoforms were indeed present in intestinal tissue (Fig. 2C). However, this analysis also detected a potentially new CDHR5 splice isoform that has a partial deletion of the first mucin-like repeat and complete deletion of the fourth repeat compared to CDHR5-L (Fig. 2C; Fig. S2B). We named this potential splice isoform CDHR5-X6 (theoretical molecular weight 76.9 kDa) in accordance with the current nomenclature used for the predicted CDHR5 splice isoforms. From our immunoblot analysis of CACO-2_BBE_ cell lysates (Fig. 2A), we interpret the ∼70 kDa band as belonging to CDHR5-S, the ∼100 kDa band as potentially belonging to CDHR5-X6, and the ∼200 kDa band being CDHR5-L. Interestingly, MLR sequence of CDHR5-L is somewhat divergent between species; mice have a 3X tandem TSP-repeat sequence in which the repeats exhibit a relatively low repeat identity, while rats have a 4X TSP-tandem repeat sequence that have a repeat identity level similar to humans (Fig. S2C). In contrast, pigs have an MLR domain that more aptly fits a 2X TSP-rich tandem repeat sequence pattern of ∼70 amino acids (Fig. S2C). Despite this diversity, every species we examined had some form of MLR domain in their CDHR5 sequence. For our analysis, we focused on characterizing the functional differences between the two dominant human splice isoforms, CDHR5-L and CDHR5-S.

To compare their cellular distributions, full-length human constructs of both CDHR5-S and CDHR5-L were stably expressed as C-terminally tagged EGFP-fusion proteins in LLC-PK1-CL4 kidney epithelial cells (Fig. 2B). LLC-PK1-CL4 cells are a pig kidney proximal tubule cell line that differentiate rapidly (∼3-4 days) to establish a robust BB on their apical surface (16). We quantified the distribution of CDHR5 by measuring the EGFP signal found in the BB versus the cytosol. We observed that when overexpressed, CDHR5-S targets efficiently to apical microvilli, with very little material found in the cytosol (Fig. 2D,F). In striking contrast, a significant amount of the CDHR5-L signal appeared trapped internally with a much lower amount reaching the apical domain (Fig. 2E, F). This demonstrates that the MLR domain found specific to CDHR5-L impedes efficient targeting to apical microvilli and may also explain the accumulation of endogenous CDHR5 signal found in the subapical region of BB microvilli from both mouse intestinal tissue and our CACO-2_BBE_ cells.

### The splice isoforms of CDHR5 can form *cis*-oligomers

In notable cases, cadherins have been shown to form *cis* interactions with themselves in order to strengthen the *trans*-adhesion that they mediate between membrane systems (17,18). Given this, we explored whether CDHR5-L could form a *cis* interaction with CDHR5-S, and if so, whether this interaction could have any influence on cellular targeting. We first assessed whether there was any apparent interaction biochemically between CDHR5-L and CDHR5-S that could indicate that they formed a *cis* complex. We co-expressed differentially-tagged versions of the isolated ectodomains (EDs) of CDHR5-L and CDHR5-S in a pairwise-manner in HEK293T cells to allow them to form potential interactions while the protein was being produced and folded inside cells. We used the native signal sequences of the EDs to direct the recombinant protein to be secreted into the medium, after which we used this conditioned medium as a source for our protein pulldowns. Interestingly, we detected a robust interaction between EDs of CDHR5-L and CDHR5-S in our pulldown assay (Fig. 3A). To determine whether this observed interaction was occurring in *cis* or *trans*, we used a fluorescent bead aggregation assay previously developed to detect *trans* adhesion interactions between cadherin molecules (19). Briefly, the EDs of CDHR5-L and CDHR5-S were expressed separately as Fc-tag proteins and adhered to different colored fluorescent Protein-A beads (Fig. S3A). The coated beads were then mixed in the presence of calcium and allowed to interact on a rocking platform. As a control, we coated beads with the ED of CDHR2 which is known to form *trans*-homophilic adhesion bonds that are strong enough to mediate bead aggregation (4,20). While we detected formation of mixed-color bead aggregates of our CDHR2 control, the CDHR5 isoforms failed to interact in *trans* to mediate bead aggregation (Fig. 3B), suggesting the interaction we observed biochemically likely represents the formation of a *cis*-oligomer between the two isoforms. We questioned whether this apparent *cis* interaction between the two splice isoforms could play a role in targeting of CDHR5-L to apical microvilli. To test this, we stably co-expressed EGFP-tagged CDHR5-L along with mCherry-tagged CDHR5-S in LLC-PK1-CL4 cells. Remarkably, expression of these two splice isoforms together in cells promoted efficient targeting of CDHR5-L to apical microvilli (Fig. 3C). We quantified this by measuring the ratio of EGFP-signal found in the BB versus the cytoplasm and found that co-expression of the two splice isoforms increased the apical targeting of CDHR5-L to levels similar to CDHR5-S (Fig. 3D; S3B,C). In sum, these results support the idea that the two CDHR5 isoforms can interact to form a *cis*-heterooligomer that targets efficiently to apical microvilli.

**Figure 3.**
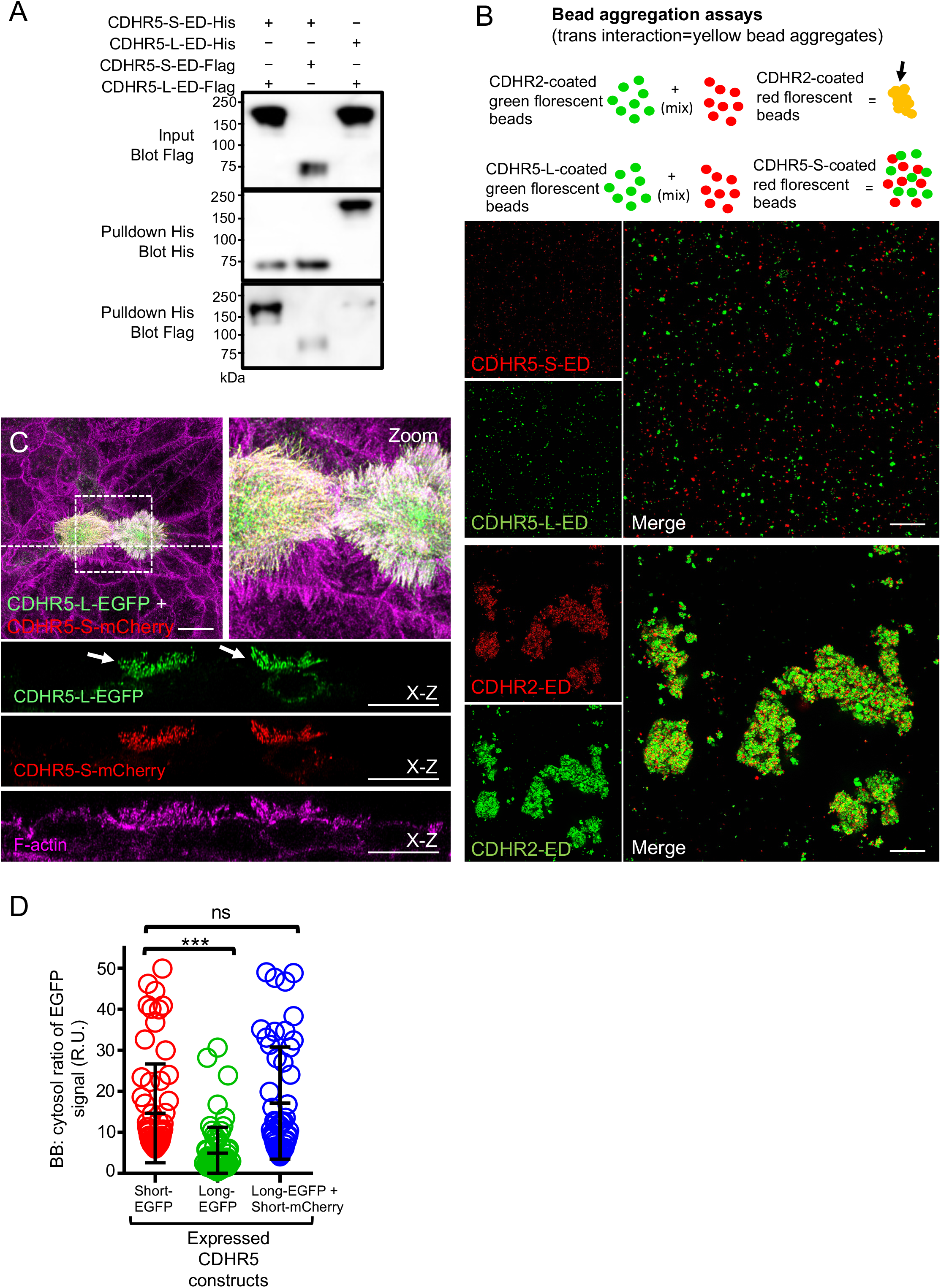
The splice isoforms of CDHR5 interact biochemically and inside kidney epithelia cells. (A) Testing for binding interactions between the EDs of the two splice isoforms of CDHR5. Differentially tagged versions of the two CDHR5 splice isoforms were co-expressed in a pairwise manner in HEK293T cells to allow for interaction and recombinant protein secretion into the medium. 6XHis-tagged constructs served as bait for pulldowns from this medium, while the Flag-tagged constructs as prey. CDHR5-S and CDHR5-L form a robust complex detected by pulldowns. (B) Bead aggregation assays to test for *trans* interactions between the two splice isoforms of CDHR5. CDHR2 is used as a control known to be able to mediate robust bead aggregation through *trans* homophilic interactions. See material and methods for experimental details. CDHR5-S and CDHR5-L do not form *trans* adhesion complex. (C) Confocal images of 4-day polarized LLC-PK1-CL4 cells stably expressing both EGFP-tagged CDHR5-L (green) and mCherry-tagged CDHR5-S (red) and stained for F-actin (magenta). Boxed region denotes area in zoomed image panel. Dashed line indicates the position where the x-z section was taken; Individual channels for the x-z section are shown below the *en face* image. Arrows in the CDHR5-L-EGFP green channel x-z section point to BB enrichment of signal. Scale bars, 10 μm. Co-expression of CDHR5-S promotes robust BB targeting of CDHR5-L. (D) Scatterplot quantification of the BB:cytosol ratios of EGFP signal from cells expressing EGFP-tagged CDHR5-S alone, EGFP-tagged CDHR5-L alone, and EGFP-tagged CDHR5-L co-expressed with mCherry-tagged CDHR5-S. Bars indicate mean and SD. R.U = relative units. ***p < 0.0001, ns=not significant, two-tailed t test.

### Overexpression of CDHR5-S promotes microvillar elongation

During the process of working with the two dominant splice isoforms of CDHR5, we observed that overexpression of CDHR5-S resulted in a dramatic change in the appearance of the BB microvilli. To explore this further, we measured the length of microvilli from cells overexpressing CDHR5-S and CDHR5-L. For comparison, we also measured the length of BB microvilli of neighboring cells that were not overexpressing these cadherins in our stable cell lines. We observed a striking increase in microvillar length specifically from cells overexpressing CDHR5-S, suggesting that this cadherin may help support microvillar elongation upon reaching the apical surface (Fig. 4A,C). In many cases for cells overexpressing high amounts of CDHR5-S, the apical domain often appeared to herniate outwards with long microvilli splaying from the herniated membrane (Fig. S4A). This could also be easily seen using scanning electron microscopy to observe the general surface features of the cell monolayer from our CDHR5-S stable cell line versus wild-type control cells (Fig. S4B,C). In contrast, overexpression of CDHR5-L did not have a similar effect on promoting elongation of microvilli (Fig. 4C). Taken together, this demonstrates that overexpression of CDHR5-S results in the elongation of BB microvilli, implicating CDHR5 in playing a key role in microvillar growth in transporting epithelia.

**Figure 4.**
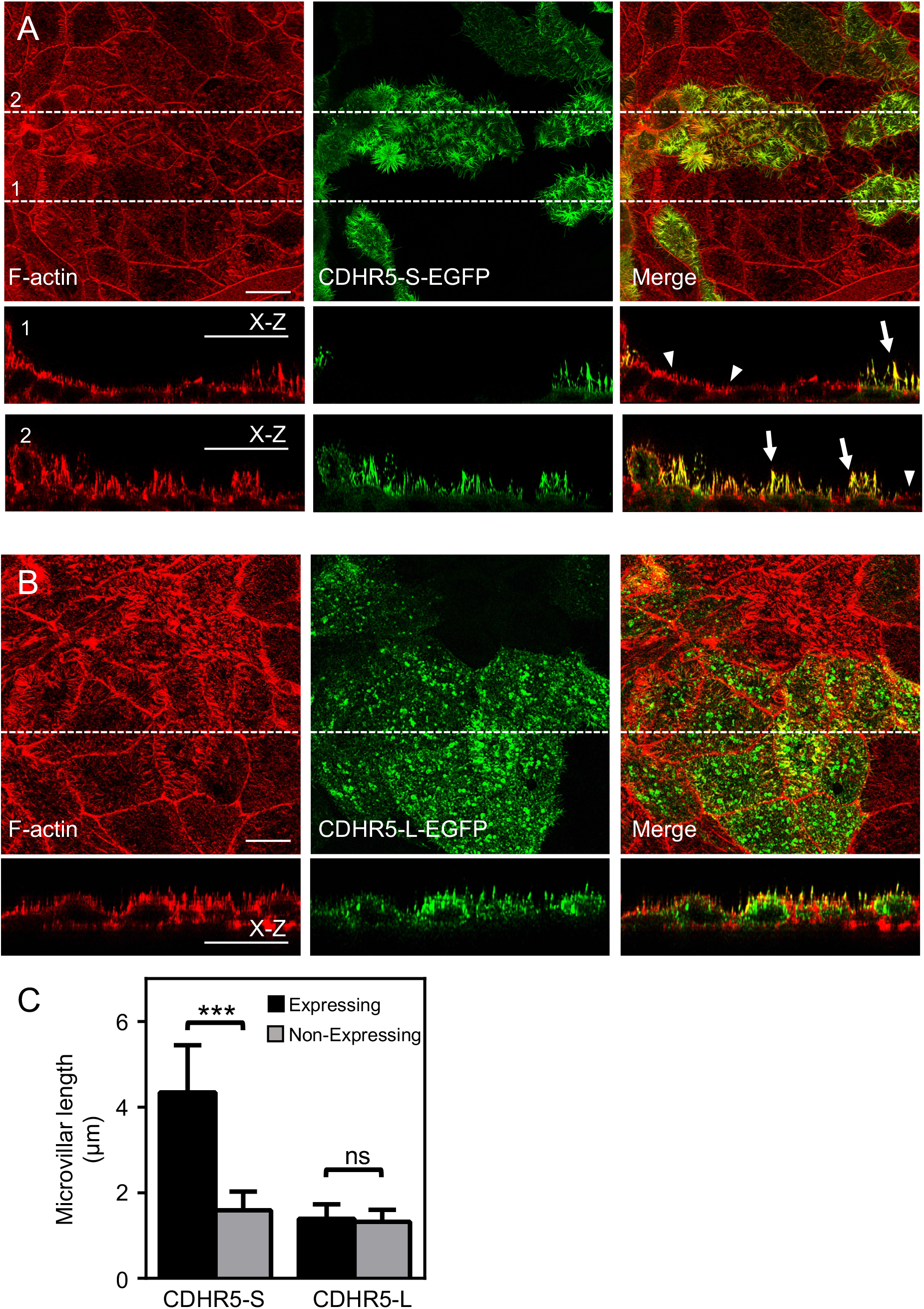
Overexpression of CDHR5-S promotes elongation of microvilli. (A+B) Confocal images of 4-day polarized LLC-PK1-CL4 cells stably expressing either EGFP-tagged CDHR5-S or CDHR5-L (green) and stained for F-actin (red). Boxed regions denote area in zoomed image panels. Dashed lines indicate the position where the x-z sections were taken; Individual channels for the x-z sections are shown below the *en face* images. Two examples of x-z sections are shown for the CDHR5-S-EGFP expressing monolayer. Arrows in the CDHR5-S-EGFP x-z section merge channel point to long microvilli, while arrowheads denote non-expressing cells that have normal, shorter microvilli. Scale bars, 10 μm. Overexpression of CDHR5-S promotes the elongation of BB microvilli. (C) Quantification of microvillar length from cells overexpressing CDHR5 isoforms compared to non-expressing cells. Bars indicate mean ± SD. Measurements: CDHR5-S expressing, N=91 microvilli, CDHR5-S non-expressing, N=58 microvilli. CDHR5-L expressing, N=70 microvilli, CDHR5-L non-expressing, N=83 microvilli.

### Different sequence features impact apical targeting of the CDHR5 isoforms

We performed a domain analysis of the two CDHR5 splice isoforms in order to further shed light on the sequence determinants that guide apical targeting of this cadherin. We first explored whether the adhesive capacity of CDHR5-S was required for its proper apical trafficking. Towards this, we stably expressed a variant of CDHR5-S lacking its first extracellular cadherin (EC) repeat (Fig. 5A; CDHR5-S_ΔEC1_) which was previously shown to be strictly required to mediate *trans*-adhesion bond formation with CDHR2 (6). Remarkably, deletion of EC1 from CDHR5-S essentially abolished apical targeting, resulting in the formation of large prominent intracellular puncta of EGFP signal (Fig. 5B; arrows). We measured the volume distribution of these puncta to compare with those seen with overexpression of CDHR5-L. On average, the puncta generated by CDHR5-S_ΔEC1_ were ∼25-fold larger than those generated by overexpressing CDHR5-L (Fig. S5A). We reasoned that removal of the entire EC1 repeat may perturb correct apical trafficking of the cadherin through other mechanisms beyond disrupting its adhesive capacity, such as removal of key glycosylation sites that may act as apical sorting signals. Therefore, we generated and tested a single point mutation in the CDHR5 EC1 repeat (CDHR5-S_R109G_) that we previously showed eliminated its *trans*-adhesion interaction with CDHR2 (4). We found that the targeting efficiency of CDHR5-S_R109G_ was indeed reduced compared to wild-type CDHR5-S (Fig. S5B-D), and that the CDHR5-S_R109G_ variant began to accumulate as puncta in the cells. These puncta, however, were much smaller in size compared to those generated by CDHR5-S_ΔEC1_. We next tested whether two reported N-glycosylation sites found in EC1 of CDHR5-S (N44 and N81) (21) were acting as important apical sorting signals that could contribute to the mistargeting of the CDHR5-S_ΔEC1_ construct. When stably expressed in our epithelial cell model, CDHR5-S_N44,81Q_ failed to target to apical microvilli (Fig. S5B,E), but was only occasionally found as large intracellular puncta. Finally, we deleted the cytoplasmic tail of CDHR5-S to determine whether it was important for apical targeting (Fig. 5A). In contrast to the EC1 domain, loss of the cytoplasmic tail of CDHR5-S had no significant impact on its ability to target to BB microvilli (Fig. 5C,F). Deletion of the cytoplasmic domain of CHDR5-S did, however, partially mute the ability of the overexpressed protein to promote microvillar elongation (Fig. 5H).

**Figure 5.**
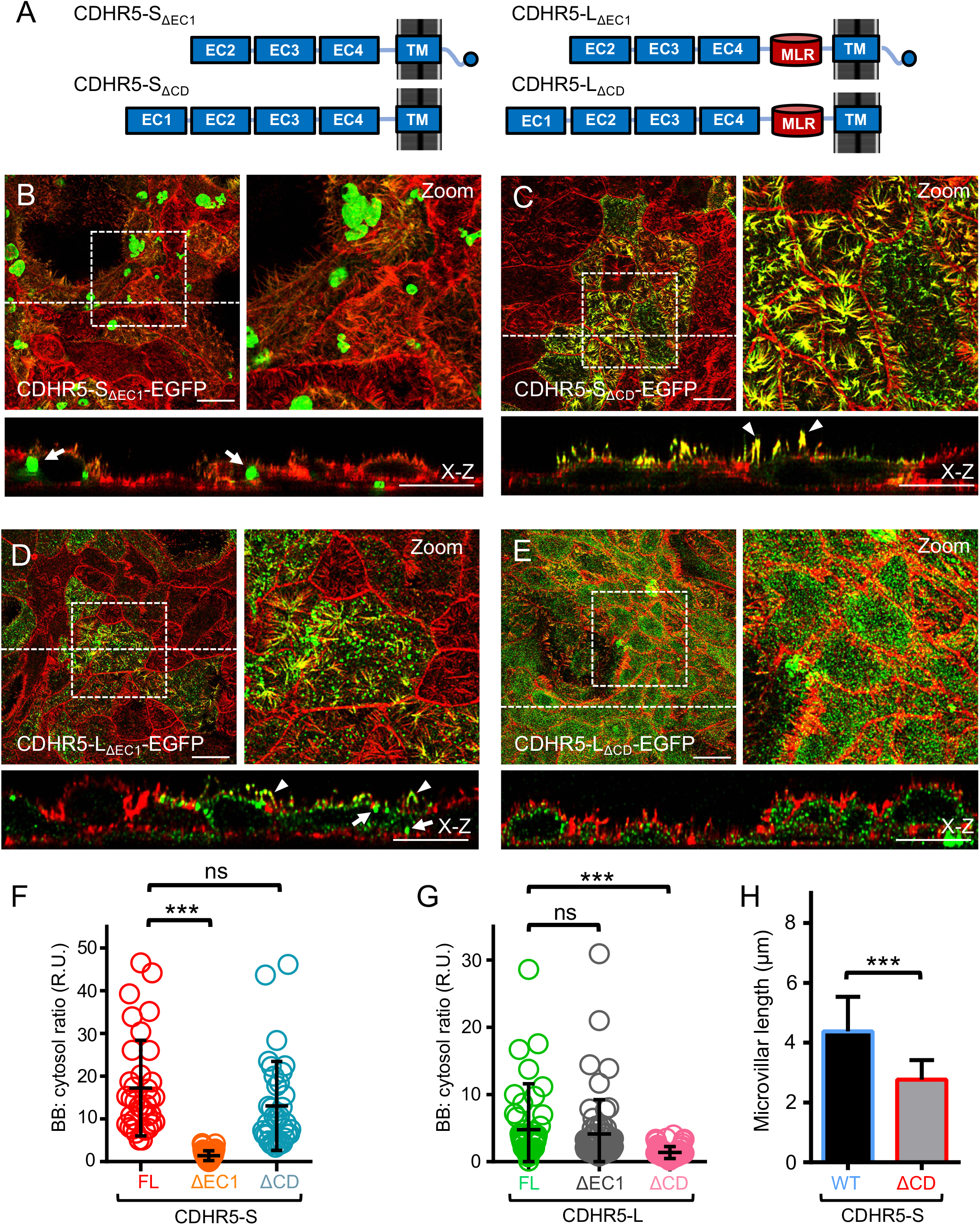
Different sequence features across the CDHR5 isoforms control their targeting properties. (A) Diagram of EGFP-fusion constructs used to explore the sequence features of the two CDHR5 splice isoforms required for apical targeting. All constructs contained an intact, native N-terminal signal sequence for correct processing by the biosynthetic pathway. (B-E) Confocal images of 4-day polarized LLC-PK1-CL4 cells stably expressing EGFP-tagged constructs tested (green) stained for F-actin (red). Boxed regions denote area in zoomed image panels. Dashed lines indicate the positions where the x-z section were taken; x-z sections are shown below each *en face* image. Arrowheads denotes localization to BB microvilli, while arrows point to signal accumulation found in subapical and cytoplasmic puncta. Scale bars, 10 μm. Different sequence features across CDHR5-S and CDHR5-L control their apical targeting. (F-G) Scatterplot quantification of the BB:cytosol ratios of EGFP signal from cells expressing the EGFP-tagged CDHR5 constructs tested. Bars indicate mean and SD. R.U = relative units. ***p < 0.0001, two-tailed t test, ns=not significant. (H) Quantification of microvillar length from cells overexpressing EGFP-tagged wild-type CDHR5-S and CDHR5-S lacking the cytoplasmic tail. Bars indicate mean ± SD. Measurements: CDHR5-S expressing, N=92 microvilli, CDHR5-S_ΔCD_ expressing, N=103 microvilli.

Surprisingly, CDHR5-L exhibited near opposite sequence-dependent targeting properties compared to CDHR5-S. Deletion of EC1 from CDHR5-L had no significant effect on its behavior; low amounts of the cadherin were still able to reach apical microvilli and only small intracellular puncta of signal were present in the cytoplasm (Fig. 5D,G). In contrast, deletion of the cytoplasmic tail resulted in a near-complete loss of apical targeting of CDHR5-L (Fig. 5E,G). We noted that in most cells however, the CDHR5-L_ΔCD_ signal did not accumulate as intracellular puncta. We next tested whether co-expressing CDHR5-S alongside our CDHR5-L variants could influence their targeting. Apical targeting of CDHR5-L_ΔEC1_ was enhanced when co-expressed with CDHR5-S (Fig. S5F), while there was only a modest improvement in BB targeting of CDHR5-L_ΔCD_ variant (Fig. S5G). In sum, these findings reveal that different sequence features across the two dominant CDHR5 splice isoforms guide BB targeting and that the cytoplasmic tail of CDHR5-L is critical for its delivery to the apical domain.

### CDHR5 interacts biochemically with E3KARP/EBP50

The cytoplasmic tail of human CDHR5 is relatively small in size (154 amino acids), is rich in glycine and proline residues with some potential poly-proline motifs, and also ends with a canonical PBM. Sequence alignment across a number of different species revealed that the C-terminal PBM is somewhat variable, conforming to either a class I or class II PBM consensus sequence (Fig. 6A). We speculated that tagging CDHR5 with a C-terminal EGFP would likely disrupt the proper function of its PBM, since a canonical PBM requires a free C-terminal hydroxyl group as part of its binding interaction (22,23). To determine whether tagging CDHR5-L with a C-terminal EGFP was interfering with its targeting ability, we expressed an untagged version of this isoform and visualized its targeting using immunofluorescence. We first confirmed that our antibody directed against human CDHR5 did not detect signal from endogenous porcine CDHR5 that could be potentially found in our LLC-PK1-CL4 cells (Fig. 6B). Strikingly, we observed that the untagged CDHR5-L had a significantly higher level of BB targeting compared to the EGFP-tagged molecule, though the levels were still below the targeting seen with CDHR5-S (Fig. 6C-E). In contrast, a similar experiment done with CDHR5-S did not detect differences between the tagged and untagged versions (data not shown), further confirming that these two CDHR5 splice isoforms exhibit different sequence-dependent targeting properties.

**Figure 6.**
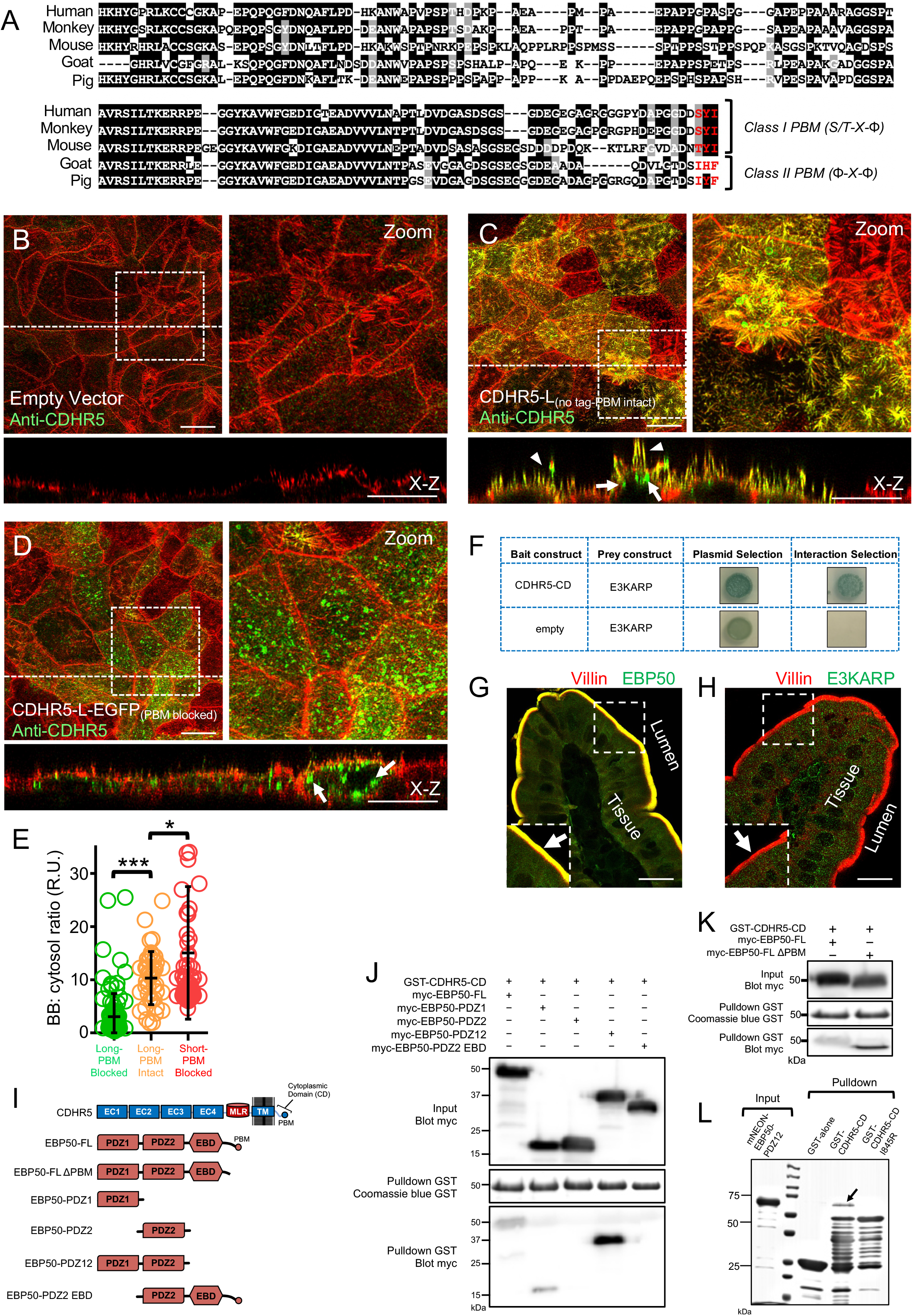
The cytoplasmic tail of CDHR5 interacts with the Ezrin-associated scaffolds E3KARP and EBP50. (A) Sequence alignment of the cytoplasmic tail of CDHR5 from Human (Homo sapiens), Rhesus Macaque (Macaca mulatta), Mouse (Mus musculus), Goat (Capra hircus), and Pig (Sus scrofa). Amino acids shaded in black are identical. Amino acids shaded in gray are similar. Amino acids colored in red denote PDZ binding motifs (PBM) X= any residue, Φ=hydrophobic residue. (B-D) Confocal images of 4-day polarized LLC-PK1-CL4 cells stably transduced with an empty-vector control, CDHR5-L with no tag (PBM intact) and EGFP-tagged CDHR5-L (PBM blocked). Monolayers have been stained for F-actin (red) and an anti-CDHR5 antibody (green) raised against the human CDHR5 sequence. Boxed regions denote area in zoomed image panels. Dashed lines indicate the positions where the x-z section were taken; x-z sections are shown below each *en face* image. Arrowheads denotes localization to BB microvilli, while arrows point to signal accumulation found in subapical and cytoplasmic puncta. Scale bars, 10 μm. CDHR5-L with an intact PBM exhibits enhanced apical targeting compared to CDHR5-L tagged with EGFP. (E) Scatterplot quantification of the BB:cytosol ratios of CDHR5 signal from cells expressing the CDHR5-L with no tag (PBM intact), EGFP-tagged CDHR5-L (PBM blocked) and EGFP-tagged CDHR5-S (PBM blocked). Bars indicate mean and SD. R.U = relative units. ***p < 0.0001, two-tailed t test. (F) Screening of a yeast-two-hybrid kidney cDNA library identified E3KARP as a binding partner for the cytoplasmic tail of CDHR5. Shown are yeast co-transformed with either the CDHR5 cytoplasmic tail bait and the recovered E3KARP prey construct or empty bait and the recovered E3KARP prey construct. Yeast were plated on a medium formulated for plasmid selection alone or a medium formulated for plasmid and interaction selection. (G+H) Confocal images of mouse duodenal tissue stained for EBP50 or E3KARP (green) and Villin (red). Arrows point to the BB. Boxed regions denote area in zoomed image inset panels. Scale bars, 10 μm. EBP50 exhibits robust targeting to BB microvilli. (I) Diagram of EBP50 constructs used to map interaction with CDHR5. (J-K) Mapping the interaction between EBP50 and the cytoplasmic tail of CDHR5. Beads coated with bacterially-expressed GST-CDHR5 CD served as bait, while COS7 cell lysates expressing myc-tagged EBP50 constructs served as pulldown material containing the prey. CD= cytoplasmic domain, FL= Full-length, PDZ12=fragment comprised of PDZ1 and PDZ2, EBD=Ezrin binding domain. The cytoplasmic tail of CDHR5 interacts with the open, active version of EBP50 through PDZ1. (L) Interaction between EBP50 and CDHR5 using purified recombinant protein from bacteria. GST-tagged purified CDHR5 proteins served as bait whereas mNEON-tagged EBP50-PDZ12 served as prey. GST alone is included as a negative control.

In order to identify potential cytoplasmic binding partners of CDHR5, we performed a yeast-two-hybrid screen of its isolated cytoplasmic tail against a kidney cDNA library and discovered the Ezrin-associated PDZ-based scaffold NHE3 kinase A regulatory protein (E3KARP; also known as NHERF-2 and SLC9A3R2) as a putative interactor (Fig. 6F). Previous work has shown that E3KARP exhibits a restricted tissue distribution, being found mostly in the epithelia of the lung and small amounts in the kidney (24). Our discovery of E3KARP as a candidate binding partner prompted us to also explore whether the closely related homolog Ezrin-radixin-moesin-binding phosphoprotein 50 (EBP50; also known as NHERF-1 and SLC9A3R1) may also interact with CDHR5. EBP50 is highly expressed in the transporting epithelia of both the kidney and small intestine, and, like E3KARP, is known to act as a scaffold to link Ezrin to a variety of transmembrane proteins found in the BB (24,25). We first confirmed that EBP50 was expressed in the intestine by staining mouse duodenal tissue sections and CACO-2_BBE_ enterocytes (Fig. 6G; S6A). In both cases, EBP50 was found highly enriched in BB microvilli. In contrast to the prominent expression of EBP50, very low amounts of E3KARP were detected in duodenal intestinal tissue and in CACO-2_BBE_ cells (Fig. 6H; Fig. S6B).

E3KARP and EBP50 have an identical domain structure, being comprised of two PDZ domains followed by an Ezrin binding domain (EBD) that terminates with a canonical class I PBM (Fig. S6C)(25). We proceeded to map the binding interactions between CDHR5 and EBP50/E3KARP using protein pulldowns. A GST-fusion protein of the cytoplasmic tail of CDHR5 was coupled to beads and used to perform pulldowns from cell lysates of COS7 cells transfected with various fragments of EBP50/E3KARP (Fig. 6I; S6D). These pulldowns revealed that the CDHR5 tail interacts with PDZ1 in EBP50 and PDZ2 in E3KARP (Fig. 6J; S6E). However, we observed that a concatenated fragment of PDZ1-PDZ2 for both EBP50 and E3KARP displayed a potentiated interaction with CDHR5 (Fig. 6J; S6E). Interestingly, our pulldowns also demonstrated that CDHR5 only weakly interacts with either of the full-length scaffolds (Fig. 6J; S6E). We speculated that this may be due to the full-length scaffolds being found in autoinhibited “closed” conformations, in which their C-terminal PBMs bind back to one of their PDZ motifs (25). It is currently proposed that EBP50/E3KARP must first interact with activated Ezrin to relieve autoinhibition before they can interact with transmembrane proteins (26,27). To test this, we deleted the C-terminal PBMs from the scaffolds to lock them in an open, active state (Fig. 6I; S6D). Consistent with this idea, we observed that the cytoplasmic tail of CDHR5 exhibited a robust interaction only with the open, active versions of both scaffolds (Fig. 6K; Fig. S6F). Finally, to validate that our observed interaction between EBP50 and CDHR5 is direct, we purified a PDZ1-PDZ2 fragment of EBP50 as a His-mNeon-tagged protein from bacteria to test with our GST-CDHR5 cytoplasmic domain. We also created a variant of the CDHR5 cytoplasmic tail in which we mutated the C-terminal residue (I845R) in order to ablate the consensus PBM which requires a hydrophobic amino acid at the C-terminal position. As expected, the PDZ1-PDZ2 fragment of EBP50 could only bind to wild-type CDHR5 cytoplasmic tail and not the CDHR5 variant lacking a canonical PBM (Fig. 6L). We conclude that CDHR5 uses its C-terminal PBM to interact directly with EBP50.

### CDHR5 interacts with E3KARP/EBP50 in cells

To further validate that EBP50/E3KARP can interact with CDHR5 in a cellular context, we took advantage of a ‘nanotrap’ pulldown approach that allows protein-protein interactions to be rapidly interrogated in live cells by assessing whether a prey protein co-localizes with a bait protein that has been forcibly targeted to the tips of filopodia using the motor activity of Myosin-10 (28). We fused various EBP50 and E3KARP constructs to mCherry to utilize as prey, while the cytoplasmic tail of CDHR5 was fused to a minimal motile form of EGFP-Myosin-10 to create a bait construct that constitutively targets to the tips of filopodia. We transfected EGFP-Myo10-CDHR5-CD along with mCherry-EBP50/E3KARP constructs in HeLa cells and plotted the correlation between bait and prey fluorescence at the tips of filopodia (Fig. 7A-F; Fig S7A-E). We observed the strongest co-localization between bait and prey when using the PDZ1-PDZ2 constructs of EBP50/E3KARP (Fig. 7D,F; Fig. S7D,E). Consistent with our conventional protein pulldown data, the observed interactions were dependent upon having EBP50/E3KARP scaffolds in the open conformation, as well as having an intact C-terminal PBM for CDHR5. In sum, our results show that the Ezrin-associated scaffolds EBP50 and E3KARP can interact with CDHR5 both biochemically and in cells.

**Figure 7.**
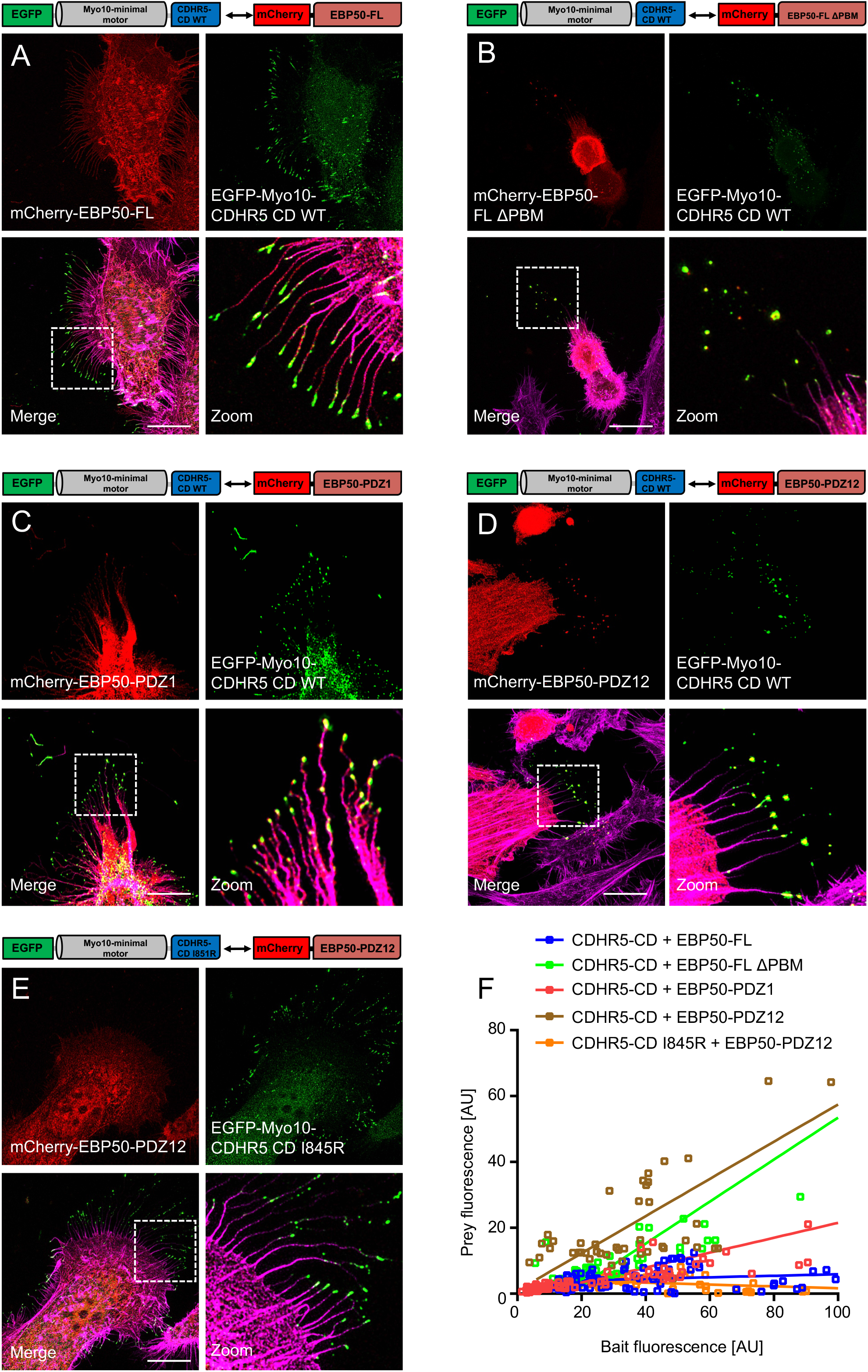
EBP50 can interact with the cytoplasmic tail of CDHR5 in a cellular context. (A-E) Confocal images of nanotrap pulldowns experiments between the cytoplasmic tail of CDHR5 and EBP50 constructs. See text for nanotrap pulldown experimental details. GFP-tagged Myo10-HMM-CDHR5 CD constructs served as bait, mCherry-tagged EBP50 constructs served as prey. Boxed regions in merge panels denote area in zoomed image panels. Scale bars,10 μm. EBP50 can interact with the cytoplasmic tail of CDHR5 in a cellular context. (F) Quantification of the correlation between bait (x-axis) and prey (y-axis) fluorescence at individual filopodia tips. Lines represent best linear fit. CD= cytoplasmic domain, FL= Full-length, PDZ12=fragment comprised of PDZ1 and PDZ2.

### CDHR5 forms a complex with Ezrin via EBP50/E3KARP in a regulated manner

The primary function of EBP50 in microvilli is to connect transmembrane proteins to Ezrin, a member of the Ezrin-Radixin-Moesin (ERM) family (25). Ezrin is known to play an essential role in the formation of microvilli by linking the plasma membrane to the underlying actin cytoskeleton (29). Ezrin is comprised of an N-terminal 4.1 protein, Ezrin, Radixin and Moesin (FERM) domain joined to a C-terminal ERM association domain (C-ERMAD) through a long α-helical coiled-coil region. In the inactive (closed) form of Ezrin, the N-terminal FERM domain binds with high affinity to the C-terminal C-ERMAD. Phosphorylation of a specific threonine residue at position 567 (T567) in the C-ERMAD, in conjunction with PI(4,5)P2 binding to the FERM domain, results in a fully active, open conformation of Ezrin (30). Active Ezrin can then bind to and activate EBP50/E3KARP using its FERM domain, while its C-terminal C-ERMAD sequence binds directly to F-actin. After mapping the CDHR5 binding site of E3KARP/EBP50, we explored whether the interaction of active Ezrin with these scaffolds would promote their open conformations to allow a ternary complex to form between CDHR5-E3KARP/EBP50-Ezrin. To test this, we generated full-length Ezrin and an Ezrin T567D phospho-mimetic mutation construct, as well as fragments that comprised the isolated FERM domain and C-ERMAD (Fig. 8A; S8A). We co-expressed these Ezrin constructs along with full-length EPB50/E3KARP in COS7 cells, and used cell lysates derived from these transfections for protein pulldowns with beads coated with GST-CDHR5 cytoplasmic tail as bait. Both the full-length and the T567D phospho-mimetic Ezrin mutant failed to initiate ternary complex formation, suggesting that the full-length inactive version of Ezrin cannot mediate this interaction and that the phospho-mimetic mutant alone does not bypass the need for PI(4,5)P2 for full activation of Ezrin (Fig.8B; S8B). In contrast, the isolated FERM domain of Ezrin appeared to activate EBP50/E3KARP to allow the ternary complex to form (Fig.8B; S8B). Importantly, the cytoplasmic domain of CDHR5 could not directly bind to Ezrin, demonstrating that complex formed was indeed a ternary interaction in which Ezrin first must activate EBP50/E3KARP, which then associates with the cytoplasmic tail of CDHR5 (Fig. 8C). Together, these data show that the available FERM domain of activated Ezrin can promote the interaction of CDHR5 with the scaffolds EBP50/E3KARP (Fig. 8D).

**Figure 8.**
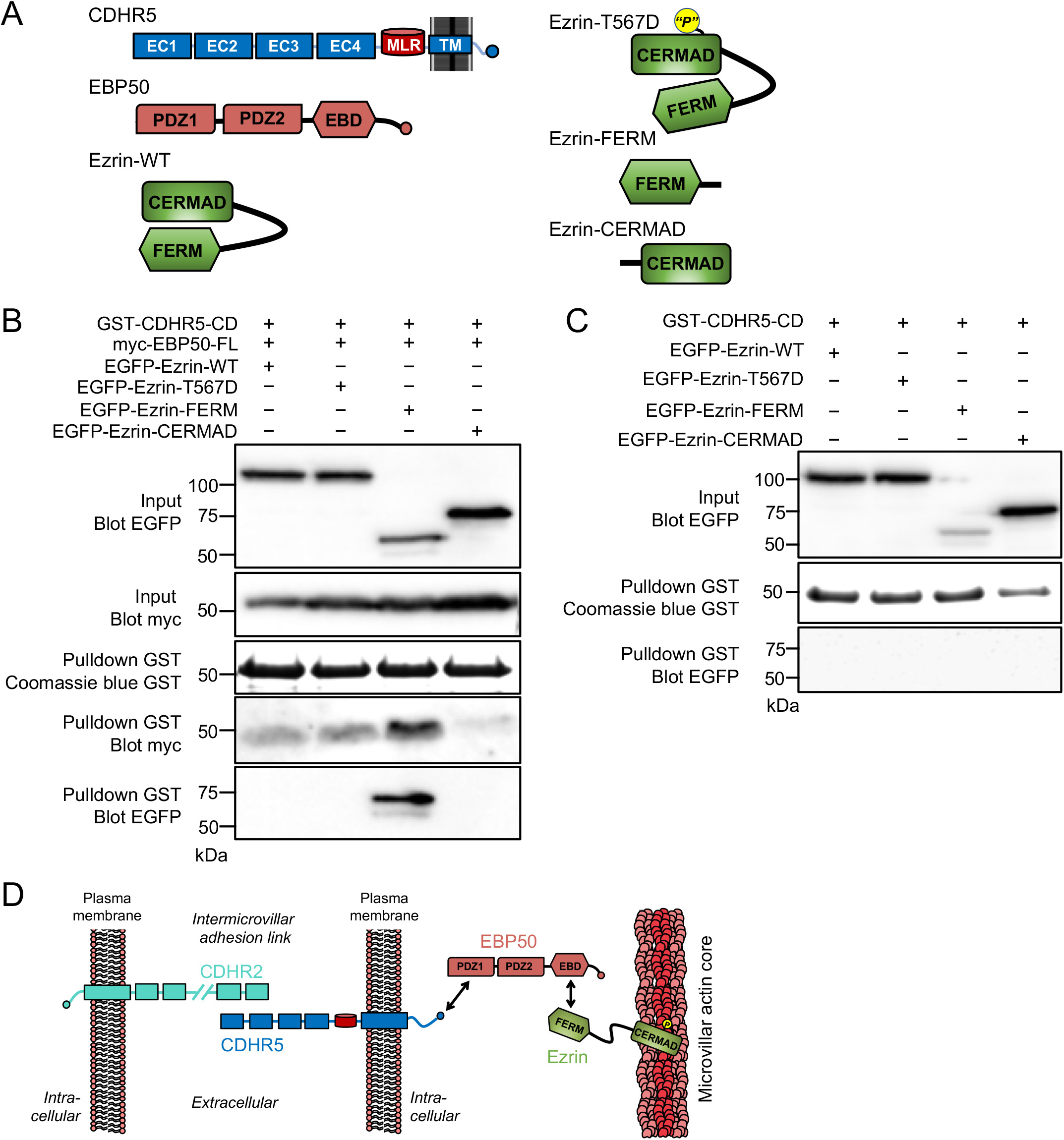
Ternary complex formation between CDHR5, EBP50, and Ezrin. (A) Cartoon schematic of the different Ezrin constructs used in pulldown assays with EBP50 and the cytoplasmic tail of CDHR5. (B) Ternary complex formation between CDHR5, EBP50 and Ezrin. Beads coated with bacterially-expressed GST-CDHR5 CD served as bait, while COS7 cell lysates expressing myc-tagged EBP50 and various EGFP-tagged Ezrin constructs served as pulldown material containing the prey. A fragment of Ezrin can activate EBP50 to allow it to interact with the cytoplasmic tail of CDHR5. (C) Pulldown analysis testing for potential interaction between CDHR5-CD and Ezrin constructs. (D) Cartoon schematic of the ternary complex formed between CDHR5, EBP50 and Ezrin.

### EBP50 is required for proper apical polarization and IMAC assembly

Given the prominent role that EBP50 plays in linking BB-resident transmembrane proteins to the actin cytoskeleton via Ezrin, we were interested to explore what the impact of loss of EBP50 would be on IMAC localization and BB assembly. We screened a panel of six independent shRNA knockdown (KD) constructs targeting EBP50 in our CACO-2_BBE_ cells and identified two that had a significant knockdown as judged by both immunostaining and immunoblot analysis (Fig. S9A,B). Consistent with previous findings in other epithelial cell systems (31,32), loss of EBP50 had a severe effect on the proper development of apical microvilli in our polarized CACO-2_BBE_ cells (Fig. 9A). Apical microvilli appeared small and rudimentary in these EBP50 KD cells, and did not cluster together indicating defective IMAC activity (Fig. 9A,B). Indeed, staining for various components of the IMAC (CDHR5, CDHR2 and USH1C) in the EBP50 KD cells revealed significant defects in their proper targeting as compared to cells transduced with a scramble control shRNA (Fig. 9C; S9B). We quantified this by measuring the ratio of signal of the IMAC components found in the apical domain compared to the total cell content. In each case, depletion of EBP50 resulted in loss of CDHR5, CDHR2, and USH1C out of BB microvilli (Fig. 9D). Immunoblot analysis of total cell lysates from these cell lines further demonstrated that the overall levels of these IMAC components were markedly reduced in the absence of EBP50 expression (Fig. S9D). We performed the complimentary experiment of knocking down CDHR5 in CACO-2_BBE_ cells to understand whether EBP50 was dependent upon this IMAC component for its proper apical targeting. Similar to loss of EBP50, KD of CDHR5 resulted in significant defects in BB assembly (Fig. 9E), with cells having only small microvilli that did not exhibit the typical clustering phenotype as seen in the scramble control cell line. In contrast however, loss of CDHR5 only had a minor impact on the efficiency of EBP50 targeting to the BB (Fig. 9E,F). We extended this analysis to look at the BB targeting efficiency of total and activated Ezrin (P-ERM; see figure legend for details) in CDHR5 KD cells. In both cases, the targeting efficiency of Ezrin did not appear to be impacted with loss of CDHR5 (Fig. S9E). Together, these studies show that proper targeting and function of the IMAC is dependent upon EBP50 in enterocytes.

**Figure 9.**
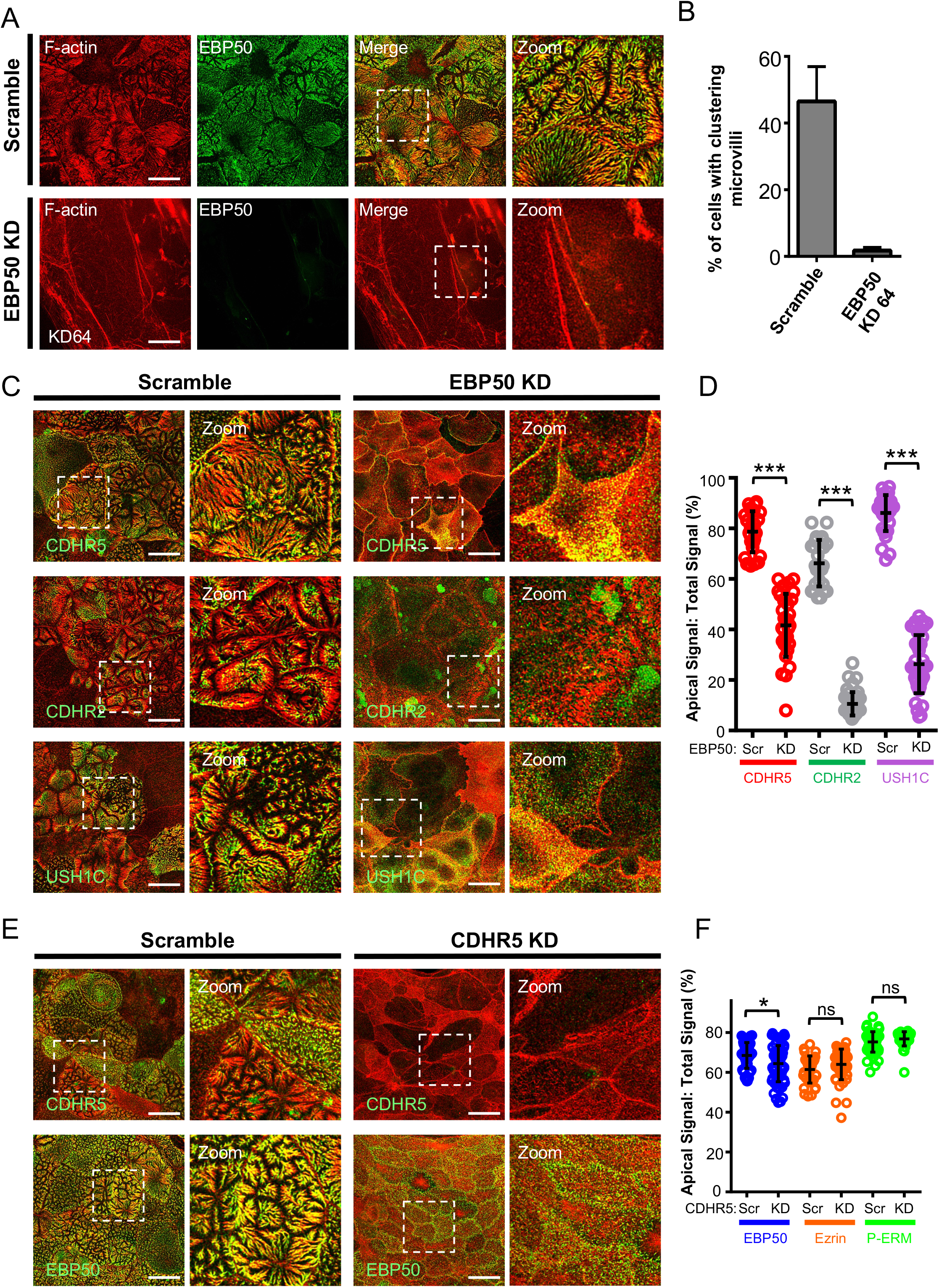
Loss of either EBP50 or CDHR5 results in severe BB assembly defects in CACO-2_BBE_ cells. (A) Confocal images of 12-day polarized CACO-2_BBE_ cells stably expressing either a scramble shRNA construct or an shRNA targeting EBP50 (KD 64), stained for EBP50 (green) and F-actin (red). Loss of EBP50 expression disrupts proper apical assembly. (B) Quantification of microvillar clustering in scramble and EBP50 KD CACO-2_BBE_ cell lines. Measurements: scramble control, n = 742 cells; EBP50 KD, n = 448 cells. Bars indicate mean ± SD. (C) Confocal images of 12-day polarized CACO-2_BBE_ cells stably expressing either a scramble shRNA construct or an shRNA targeting EBP50, stained for F-actin (red) and various IMAC components (CDHR5, CDHR2, or USH1C; green). Boxed regions denote the area in zoomed image panels. Scale bars, 10 μm. (D) Scatterplot quantification of apical/total signal ratios of the IMAC components in scramble and EBP50 KD CACO-2_BBE_ cells. Data are from stable cell lines that were independently derived twice. Each data point represents the ratio of the entire apical signal found in the x-z section compared to the total signal of the x-z section. ****p < 0.0001, two-tailed t test. Loss of EBP50 causes defects in IMAC targeting. (E) Confocal images of 12-day polarized CACO-2_BBE_ cells stably expressing either a scramble shRNA construct or an shRNA targeting CDHR5, stained for F-actin (red) and EBP50 (green). Boxed regions denote the area in zoomed image panels. Scale bars, 10 μm. (F) Scatterplot quantification of apical/total signal ratios of EBP50, Ezrin, and Phospho-ERM (P-ERM) in scramble and CDHR5 KD CACO-2_BBE_ cells. Data are from stable cell lines that were independently derived twice. Each data point represents the ratio of the entire apical signal found in the x-z section compared to the total signal of the x-z section. *p < 0.01, ns=not significant, two-tailed t test.

## DISCUSSION

Numerous groups have shown that expression of CDHR5 is almost always lost in colorectal cancer, which raised the possibility that this cadherin may function as a critical tumor suppressor in the gut (12,13,33-37). Direct overexpression of CDHR5 inhibited low-density proliferation of colorectal cancer cells in culture and reduced their tumor formation potential when implanted into nude mice (36), further suggesting that pharmacological manipulation of CDHR5 levels may be a future avenue for the treatment of colorectal cancer. Along these lines, the promising colorectal cancer drug mesalazine (5-aminosalicylic acid, 5-ASA) was shown to significantly upregulate CDHR5 expression in enterocytes, which may be part of its chemopreventive mode of action (38,39). However, recent studies of a CDHR5 KO mouse line did not reveal any direct differences in either lifespan or spontaneous tumor incidence between wild-type and KO littermates (37), leaving the nature of the potential tumor suppressing activity of CDHR5 unclear. In this study, we sought to examine the basic targeting properties of CDHR5 in epithelial cells to shed light on how this cadherin reaches the apical domain to exert its biological and potential tumor suppressor activity. We discovered that the two splice isoforms of this cadherin rely on significantly different targeting determinants to reach apical microvilli of the BB. Our results support the idea that the two splice isoforms can form a lateral ‘*cis*’ heterooligomer that is competent to target to apical microvilli and support microvillar growth.

In a somewhat unexpected parallel between divergent tissue systems, intermicrovillar adhesion links that connect and control BB microvilli organization in the gut and kidney bear a striking resemblance to the tip-link filaments connecting the staircase-like arrangement of stereocilia together in the inner ear that mediate hearing (1). This resemblance extends to not only the cadherins themselves, but also a majority of their cytoplasmic binding partners (4,6,7,40,41). Stereocilia tip-links are comprised of Cadherin-23 (CDH23) and protocadherin-15 (PCDH15) that interact in *trans* to form the tip-link filament that connects the tip of one stereocilia to the side of its adjacent taller neighbor (42). The emerging structural view of inner ear tip-links is that they are formed by a 2:2 complex of CDH23 and PCDH15, in which a *cis*-homodimer of CDH23 forms the upper portion of the tip-link and a PCDH15 *cis*-homodimer forms the lower portion (42-44). Sequence and structural analyses suggest that CDHR5 found in intermicrovillar adhesion links represents the ‘functional equivalent’ of PCDH15 found in inner ear tip-links (45). Detailed structural and biophysical studies showed that PCDH15 dimerizes by forming a double-helical assembly of two molecules, mediated by two independent *cis*-dimerization interfaces involving the EC2-EC3 domain region and a unique membrane-adjacent domain (MAD)(43,44). *Cis*-dimerization of the stereocilia cadherins is thought to have an important impact on the biology of the tip-link, by not only increasing the effective adhesive strength through formation of a dimeric *trans* interface, but also the nature of the helical configuration of the *cis*-dimers suggests the potential for elasticity through helix winding and unwinding (43). Both of these properties may be important to allow the tip-link to remain intact under the forces endured during the mechanotransduction mechanism of hearing. It is likely that intermicrovillar adhesion links also encounter significant forces due to shear flow of contents moving through the intestine and kidney, so having a similar cadherin conformation may be necessary to maintain intermicrovillar adhesion link integrity (1). It is interesting to note that the MAD dimerization sequence of PCDH15 is in an identical structural position compared to the MLR domain of CDHR5, being found just outside the cell membrane. Besides their location and apparent roles in *cis* oligomerization, these two sequences do not display any other overt homology. While the PCDH15 MAD sequence adopts a ferredoxin-like fold (44), the CDHR5 MLR domain clearly resembles a typical mucin tandem repeat sequence that is seen across members of the secreted and transmembrane mucin family. The CDHR5 MLR domain is likely subject to heavy O-glycosylation (10) and therefore may also contribute towards the formation of the dense glycocalyx barrier associated with the BB membrane. Furthermore, creating a potential *cis*-heterodimer of CDHR5-L and CDHR5-S in which the two ectodomains are different lengths may allow for the creation of different architectures of intermicrovillar adhesion links. We note that in our early high-resolution images of intermicrovillar adhesion links from both CACO-2_BBE_ cells and mouse enterocytes, we could easily see examples in which three intermicrovillar adhesion links originating from three neighboring microvilli would fuse centrally to couple the three microvilli together (4). This bifurcated intermicrovillar adhesion link geometry may contribute to the correct hexagonal packing of microvilli during BB assembly, as well as play a direct role in barrier function by more effectively blocking access to the space in between microvilli. Going forward, it will be important to further investigate the structure of CDHR5 with the goal of understanding the role of *cis*-oligomerization in its biology.

Our results also provide evidence that CDHR5 has a role in microvillar elongation in addition to its role in organizing the BB. Similar to our observation with CDHR5, modulation of the Drosophila melanogaster cadherin Cad99C was previously observed to control microvillar length in the fly system (46,47). Cad99C is found highly enriched in the microvilli of follicle cells that are in contact with the Drosophila oocytes, and represents the fly orthologue of PCDH15 (46). Flies depleted of Cad99C had short microvilli on their follicle cells, whereas Cad99C overexpression promoted pronounced elongation of microvilli. In combination with our study here, these findings highlight a potential conserved function of protrusion-associated cadherins in microvillar growth. The molecular basis of this observation may relate to the recent discovery that mucin biopolymers and long-chain polysaccharides found within the cell glycocalyx can generate entropic forces that promote the formation of *de novo* finger-like membrane extensions from the cell surface (48). Whether the levels of glycosylation of the ectodomains of Cad99C and CDHR5 correlates with their ability to promote microvillar elongation will be an interesting future question to address.

We further identified E3KARP/EBP50 as potential binding partners for the cytoplasmic tail of CDHR5 in our study. Within the gut, EBP50 is known to play an important role in linking transmembrane proteins to the actin cytoskeleton through Ezrin. Unless prevented, membrane surface tension would normally promote the coalescence and fusion of adjacent BB microvilli together in order to achieve an energetic minimum (49). By indirectly crosslinking the actin core to the overlying membrane, EBP50 counteracts this tendency. Indeed, while the EBP50 null mouse was found to be viable, it exhibited shortened malformed microvilli (32) that were comparable to the stunted microvilli seen in unviable embryos of ezrin null mice (50). In a similar manner, the other IMAC protocadherin CDHR2 is indirectly cross-linked to the actin cytoskeleton through two scaffolds, USH1C and ANKS4B, that interact with the actin binding protein Myo7b (4,6). Deletion of CDHR2 from mice also leads to shorter BB microvilli (8). More recently, EBP50 was identified as a binding partner for the cell adhesion molecule Transmembrane and Immunoglobulin Domain–containing protein 1 (TMIGD1) that is found enriched at the proximal base region of enterocyte microvilli (51). TMIGD1 was shown to be able to mediate trans-homophilic adhesion, and was proposed to form another set of potential ‘intermicrovillar adhesion links’ that are located more towards the base of microvilli. The importance of these linkages is highlighted by the fact that loss of TMIGD1 from mice results in severe BB assembly defects that mimic the loss of the IMAC (51). This again draws parallels to the formation of inner ear hair bundles that rely on different types of linkages that connect stereocilia together during hair bundle assembly. Understanding the developmental timing, molecular binding partners, and relative contribution of the different types of linkages that connect microvilli together in the BB is a clear future goal.

## MATERIAL AND METHODS

### Molecular Biology

The human cDNA constructs of CDHR2 (NM_001171976.2;UniProtKB-Q9BYE9), CDHR5-L (NM_021924.5;UniProtKB-Q9HBB8-1), CDHR5-S (NM_031264.5;UniProtKB-Q9HBB8-2), EBP50 (NM_004252.5;UniProtKB-O14745-1), E3KARP (NM_001130012.3;UniProtKB-Q15599-1), and Ezrin (NM_003379.5;UniProtKB-P15311) were used in this study. All fragment and truncation constructs including CDHR5-CD (aa 697–845), CDHR5-S_ΔEC1_ (aa 1-651, Δ71-124), CDHR5-S_ΔCD_ (aa 1–501), CDHR5-L_ΔEC1_ (aa 1-845, Δ71-124), CDHR5-L_ΔCD_ (aa 1–695), EBP50-PDZ1 (aa 1–148), EBP50-PDZ2 (aa 149–298), EBP50-PDZ12 (aa 1–298), EBP50-PDZ2 EBD (aa 149–358), EBP50-FL ΔPBM (aa 1–353), E3KARP-PDZ1 (aa 1–149), E3KARP-PDZ2 (aa 150–284), E3KARP -PDZ12 (aa 1–284), E3KARP-PDZ2 EBD (aa 150–337), E3KARP-FL ΔPBM (aa 1–329), Ezrin-WT (aa 1–586), Ezrin-NERMAD (aa 1–297), Ezrin-CERMAD (aa 461–586), were generated by PCR and TOPO cloned into the pCR8 Gateway Entry vector (Invitrogen). The Quikchange site-directed mutagenesis kit (Aligent) was used to generate CDHR5-S_R109G_, CDHR5-S_N44,81Q_, CDHR5-CD I845R, and Ezrin-T567D mutants within their entry vectors. All entry vectors clones were verified by DNA sequencing. Entry clones for targeting and interaction assessment in mammalian cell culture lines were shuttled into the appropriate destination vectors including: pEGFP-N3 (Clontech), pmCherry-C1 (Clontech), and pLVX-N3-EGFP (Clontech) that had been Gateway-adapted using the Gateway Vector Conversion kit (Invitrogen). Vectors used for expression of recombinant protein for pulldown assays including His-mNEON tagged, FLAG tagged, myc tagged, and FC tagged, and EGFP-tagged proteins in this study have been previously described (4,6,40). Knockdown studies utilized a non-targeting scramble control shRNA (Addgene; plasmid #46896), CDHR5 KD shRNA (TRC clone TRCN000054168; KD68), and EBP50 KD shRNA clones (TRCN0000043733; KD33), (TRCN0000043734; KD34), (TRCN0000043735; KD35), (TRCN0000043737; KD37), (TRCN0000440444; KD44) and (TRCN0000437164; KD64) sequences that were expressed from the pLKO.1 plasmid. The pcDNA3.1-MYO10-HMM-Nanotrap system was a gift from Thomas Friedman (Addgene plasmid # 87255; http://n2t.net/addgene: 87255; RRID: Addgene_87255). Flagged-tagged Human EBP50-FL (aa 1-358), EBP50-PDZ1(aa 1-148), EBP50-PDZ2 (aa 148-298), and EBP50-PDZ12 (aa 1-298) cloned into the pCMV-2-FLAG vector were gifts from Maria-Magdalena Georgescu (Addgene plasmids #28291-28297).

### Human Intestinal cDNA library PCR screening

Primers designed to flank the MLR of human CDHR5 were used for PCR reactions with Human Small Intestine QUICK-Clone cDNA (Clontech) as the template. PCR reactions were set up as recommended by the manufacturer. PCR products were then recovered and cloned using the pCR8-GW-TOPO system (Invitrogen) and sent for sequencing.

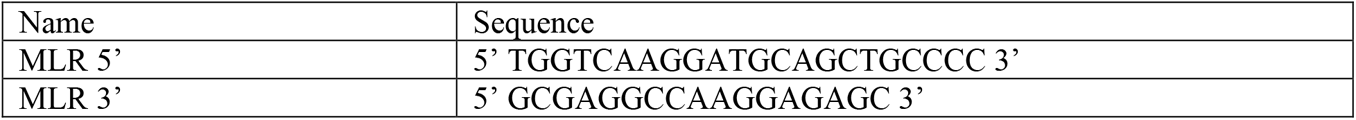

### Yeast two-hybrid screening

Yeast-two-hybrid library screening was performed using the MATCHMAKER system (Takara) with a human kidney yeast-two-hybrid cDNA library. Library-adapted cDNAs were expressed from the pACT2 vector in the *MAT*α Y187 yeast strain, while the CDHR5-CD bait construct was expressed from the pGBKT7 vector in the *MAT*a AH109 strain. Library screening, including the use of controls, was performed using standard methods outlined by the MATCHMAKER system.

### Cell culture and stable cell line generation

LLC-PK1-CL4, CACO-2_BBE_, HeLa, HEK293T and HEK293FT cells were grown in a 5% CO2 humidified incubator at 37°C in DMEM supplemented with high glucose and 2 mM L-glutamine. CACO-2_BBE_ cells were cultured in DMEM with 20% fetal bovine serum (FBS) and LLC-PK1-CL4, HeLa, HEK293T, HEK293FT cells in DMEM with 10% FBS. HEK293FT cells (10 cm dish at 80% confluency) were used for lentivirus production by co-transfecting 6 μg of Lentiviral expression plasmid with 4 μg psPAX2 packaging plasmid and 0.8 μg pMD2.G envelope plasmid using the polyethylenimine transfection reagent (Polysciences). Cells with transfection medium were exchanged with fresh medium 12 hours after transfection. Cells were returned to the 37°C incubator with 5% CO2 to allow for lentiviral production into the medium. Two days later, supernatant containing secreted lentiviral particles was collected, filtered with a 0.45 μm syringe filter and concentrated using Lenti-X Concentrator (Clontech) and stored at -80°C for further use. To transduce LLC-PK1-CL4 and CACO-2_BBE_ cells with lentivirus, cells were seeded and grown to 80% confluency in a T25 flask, after which the medium was supplemented with 8 μg/ml polybrene and ∼300μl of concentrated lentivirus was added. Cells were then incubated overnight with this transduction medium. For plasmid transfections, LLC-PK1-CL4 cells were first seeded and grown to 80% confluency in a T25 flask. Plasmids were mixed with Lipofectamine 2000 (Invitrogen) according to the manufacturer’s instructions to create a transfection mixture. This transfection mixture was added to the culture medium of cells and allowed to incubate for 3 hours. Fresh medium was then swapped in and cells were allowed to recover for 24 hours. To select for stable cell lines after viral transduction or plasmid transfection, T25 flasks containing the transduced/transfected cells were reseeded to 10 cm dishes and grown for 3 days in the absence of antibiotics. Next, cells were reseeded into T182 flasks with medium containing 50 μg/ml of puromycin for viral transductions or 1 mg/ml of G418 for plasmid transfections. Cells were then continuously grown for numerous (∼10) passages to select for stable integration of DNA.

### Recombinant protein purification from E. coli

For pulldown assays using the GST fusion of the CD of CDHR5, the pGEX-4T3-CDHR5 CD construct was transformed into BL21(DE3) bacteria (Thermo Fisher Scientific), expressed, and purified using GSH resin using standard conditions. For recombinant production of EBP50, the pNCS-NEON-EBP50 PDZ1PDZ2 construct was expressed in BL21(DE3) bacteria. Bacteria were grown to a O.D=1 at 37 °C, after which the temperature was lowered to 16 °C and expression induced for 8 hrs using ITPG (Invitrogen). Cells were then collected by centrifugation at 4000 x *g* for 20 minutes and purified using Ni–NTA–agarose (Qiagen) using standard conditions. Briefly, cells were lysed using cold lysis buffer (50 mMNaPO4, pH 8.0, 300 mM NaCl, 10 mM imidazole, 1 mg/ml lysozyme, 1 mM PMSF) along with sonication. The cell lysate was then centrifuged at 15,000 x g for 1 hr. The soluble fraction of the lysate was incubated with Ni-NTA agarose resin (Qiagen) on a rocking platform for 1 hr at 4°C. The lysate–Ni–NTA resin mixture was applied to a column, washed with Wash Buffer (50 mM NaPO4, pH 8.0, 300 mM NaCl, 25 mM imidazole) and the His fusion protein eluted with Elution Buffer (50 mM NaPO4, pH 8.0, 300 mM NaCl, 250 mM imidazole). Eluates were analyzed by SDS-PAGE and those fractions containing His-NEON-EBP50 PDZ1PDZ2 were pooled, dialyzed, and utilized for protein pulldown assays.

### HEK293T Protein Production, Bead Aggregation Assays, and Pulldown Assays

Recombinant ectodomain proteins were generated by transfection of HEK293T cell cultures using Lipofectamine 2000 (Invitrogen) according to the manufacturer’s protocol. The medium containing the secreted proteins was collected, filtered, and concentrated using Ultracel concentrators (Millipore). Media containing the recombinant ED proteins were then used directly for either bead aggregation assays or for protein pulldown assays. Bead aggregation assays were performed as previously described (20). Protein pulldowns were performed as previously described (4).

### Confocal Microscopy

CACO-2_BBE_, LLC-PK1-CL4, HeLa cells seeded on coverslips were fixed in 4% paraformaldehyde (Electron Microscopy Sciences) in PBS for 15 min at RT, washed with PBS, and permeabilized with 0.1% Triton X-100 in PBS for 7 min. After fixation, cells were washed four times with PBS and blocked overnight with 5% BSA at 4°C. Cells were stained for 1 hr at room temperature using primary antibody for anti-CDHR5 (1:200; Sigma catalog no. HPA009081), anti-CDHR2 (1:75; Sigma catalog no. HPA012569), anti-USH1C (1:70; Sigma catalog no. HPA027398), anti-E3KARP (1:200; Sigma catalog no. HPA001672), anti-EBP50 (1:200; Sigma catalog no. HPA027247), anti-Ezrin (1:200; Cell Signaling catalog no. 3145), anti-P-ERM (1:200; Cell Signaling catalog no. 3726) or anti-GFP (1:200; Aves Labs catalog no. GFP1020). Cells were then washed three times with PBS and incubated with Alexa Fluor 488 goat anti-rabbit or Alexa Fluor 488 goat anti-chicken (where appropriate) and Alexa Fluor 568 phalloidin diluted 1:200 in PBS for 1 hr at RT. Cells were washed four times with PBS and coverslips were mounted using ProLong Diamond Anti-fade reagent (Invitrogen). Paraffin-embedded intestinal tissue sections from mice were prepared as previously described (4). Cells and tissue sections were imaged using a Leica TCS SP8 laser-scanning confocal microscope equipped with HyVolution deconvolution software.

### Electron microscopy

LLC-PK1-CL4 cells were seeded into 0.4-μm collagen coated 12-mm Transwell-COL inserts (Corning) and allowed to polarize for 4 days. Samples were washed once with warm SEM buffer (100mM sucrose and 100mM Na-phosphate buffer, pH 7.4) and fixed overnight at 4°C with 3% glutaraldehyde in SEM buffer. Samples were washed with SEM buffer followed by incubation with 1% OsO4 in SEM buffer on ice for 1 h, and subsequently washed with SEM buffer. Samples were dehydrated in a graded ethanol series, dried, mounted on aluminum stubs, and coated with gold/platinum using a sputter coater. Imaging was performed using a JEOL JSM-7500F field emission scanning electron microscope operated in high vacuum mode with an accelerating voltage of 0.5-2 kV. All SEM reagents were purchased from Electron Microscopy Sciences.

### Statistical analysis

All graphs were generated and statistical analyses performed using Prism version 6 (GraphPad). For all figures, error bars represent S.D. Unpaired t tests were employed to determine statistical significance between reported values. Statistical details of individual experiments can be found in figure legends (***, p < 0.0001; **, p< 0.001; *, p <0.01).

## FIGURE LEGENDS

**Figure S1.**
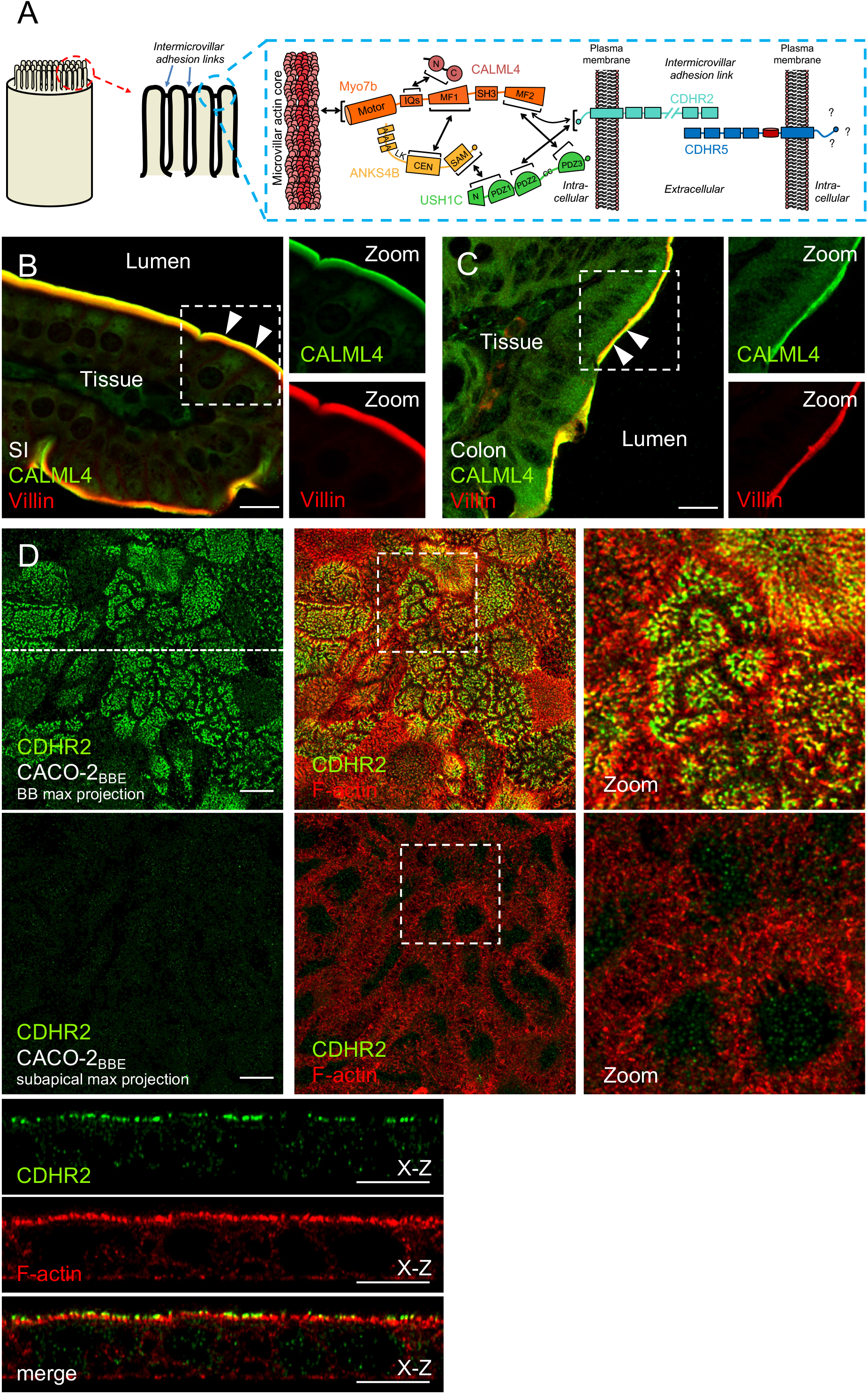
Targeting properties of other IMAC components in intestinal tissue and CACO-2_BBE_ cells. (A) Cartoon showing the location of intermicrovillar adhesion links found at the tips of BB microvilli and the interactome of the IMAC. (B+C) Confocal images of mouse small intestine (SI) and colon tissue stained for CALML4 (green) and Villin (red). Arrowheads denotes localization to the tips of BB microvilli. Boxed regions denote area in zoomed image panels. Zoom images are shown as individual channels. Scale bars, 10 μm. Unlike CDHR5, CALML4 does not exhibit distinct enrichment in puncta below the BB of intestinal tissue. (D) Confocal images of 12-day polarized CACO-2_BBE_ cells stained for endogenous CDHR2 (green) and F-actin (red). Upper panel series show a maximum projection through the full height of the BB, while lower panels show a maximum projection through the same monolayer excluding the apical BB. Boxed regions denote area in zoomed image panels. Dashed line in the BB maximum projection indicate the position where the *x-z* section was taken; Individual channels for the *x-z* section are shown below the *en face* image. Scale bars, 10 μm. Unlike CDHR5, CDHR2 does not exhibit distinct enrichment in puncta below the BB of CACO-2_BBE_ cells.

**Figure S2.**
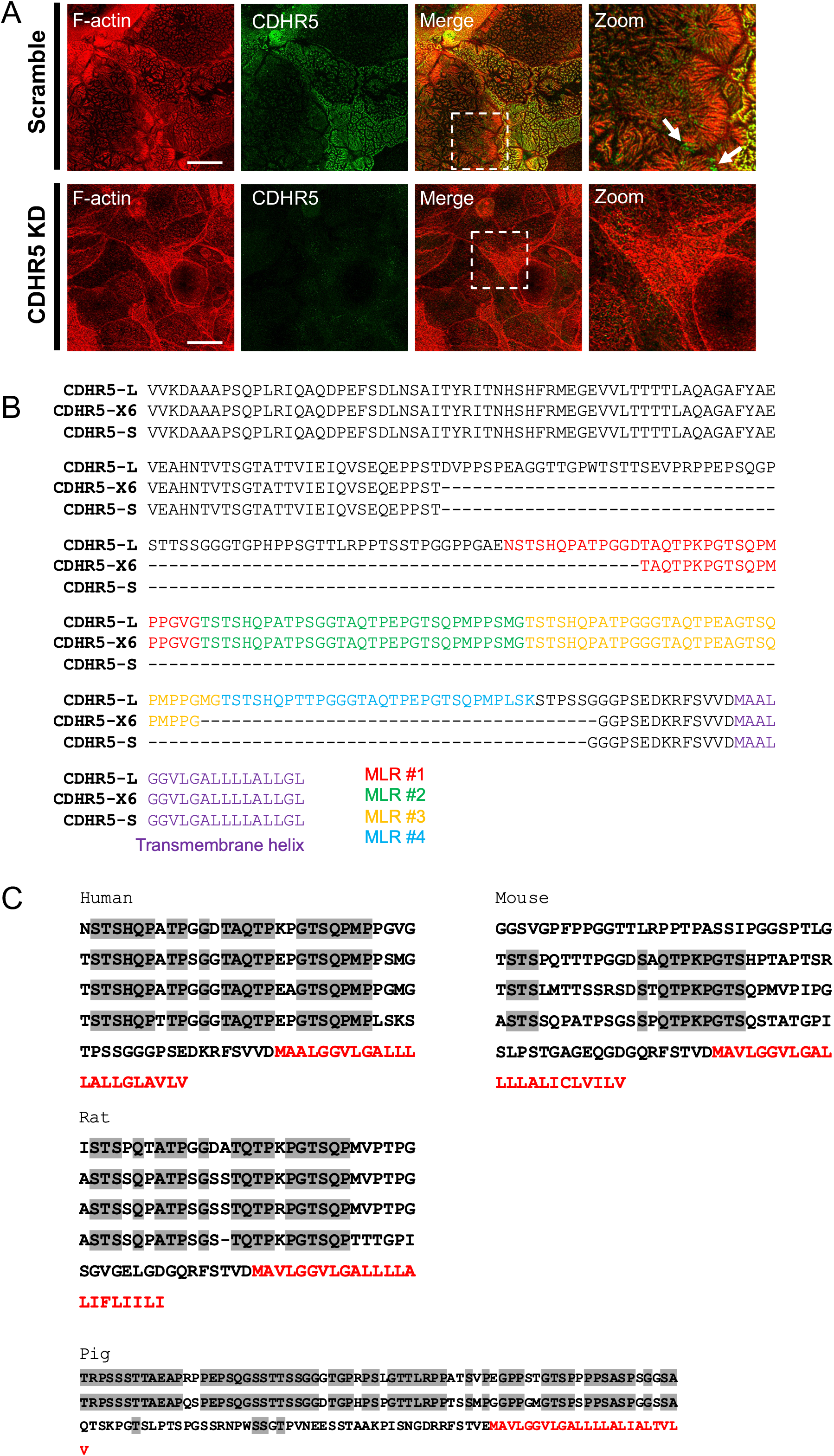
CDHR5 antibody validation and sequence analysis of the MLR domain. (A) Confocal images of 12-day polarized CACO-2_BBE_ cells stably expressing either a scramble shRNA construct or an shRNA targeting CDHR5, stained for CDHR5 (green) and F-actin (red). Arrows point to internally trapped material. Images show that our CDHR5 antibody does not exhibit off-targeting labeling. (C) Sequence alignment of the MLR domain between CDHR5-L, CDHR5-S and the identified isoform CDHR5-X6. CDHR5-X6 lacks the fourth mucin-like repeat and has a partial deletion of the first repeat. The Mucin-like repeats, as well as the transmembrane helix, are color-coded as shown. (C) Internal sequence alignments of the MLR tandem repeats found in CDHR5 across selected species (human, mouse, rat and pig). Grey highlighted residues indicate identity across the mucin-like tandem repeats within each species. Residues in red denote the predicted transmembrane sequences for each CDHR5 protein.

**Figure S3.**
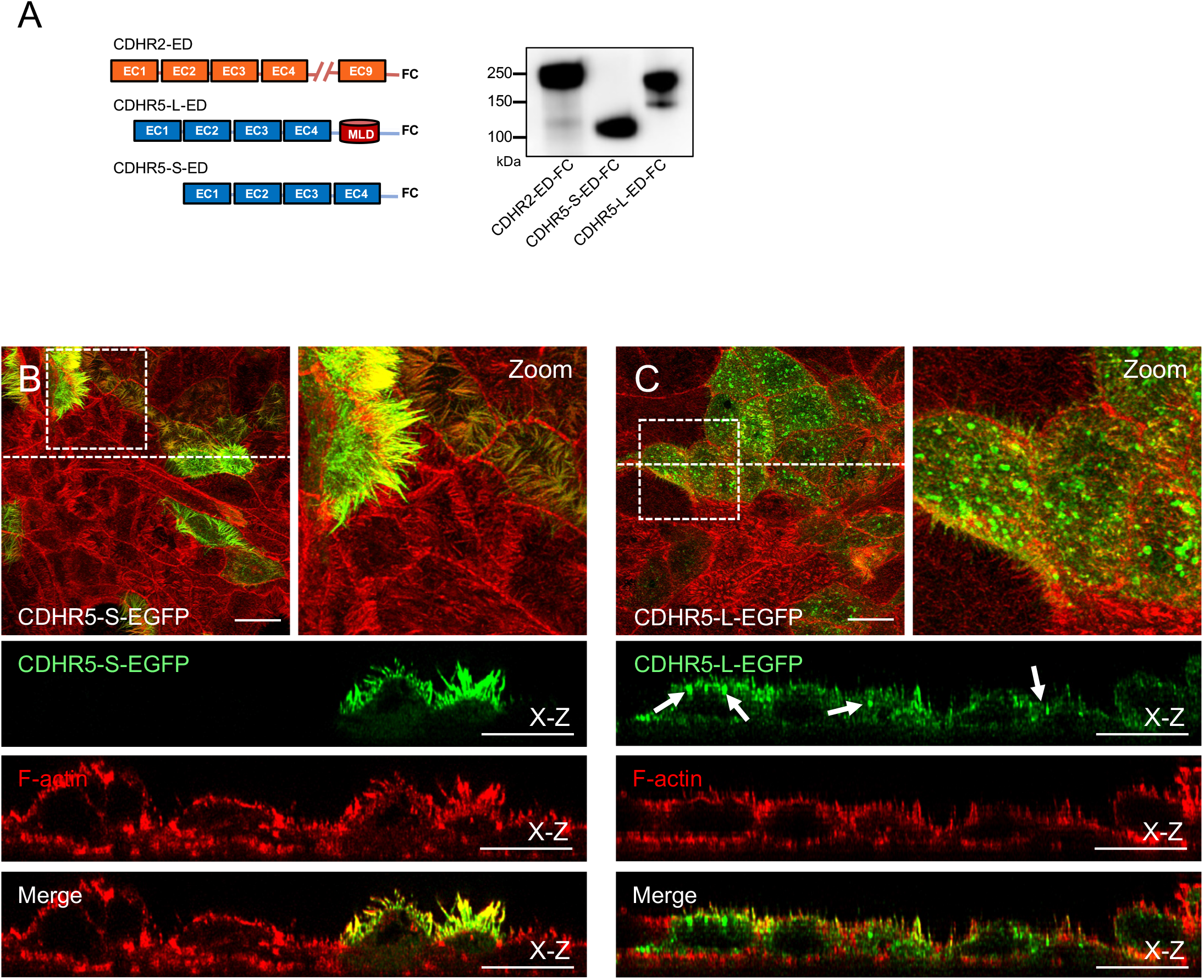
Reagents and control experiments for testing whether the two splice isoforms of CDHR5 interact biochemically and inside kidney epithelia cells. (A) Domain diagram and immunoblot analysis of constructs used to label florescent beads for bead aggregation assays. Immunoblot was performed with an anti-Fc antibody. (B+C) Confocal images of 4-day polarized LLC-PK1-CL4 cells stably expressing either EGFP-tagged CDHR5-L or CDHR5-S (green) and stained for F-actin (red). Boxed regions denote area in zoomed image panels. Dashed lines indicate the position where the x-z sections were taken; Individual channels for the x-z sections are shown below the *en face* images Arrows in the CDHR5-L-EGFP green channel x-z section point to subapical and cytoplasmic puncta of signal. Scale bars, 10 μm. In the absence of co-expression of CDHR5-S, CDHR5-L exhibits poor targeting to BB microvilli.

**Figure S4.**
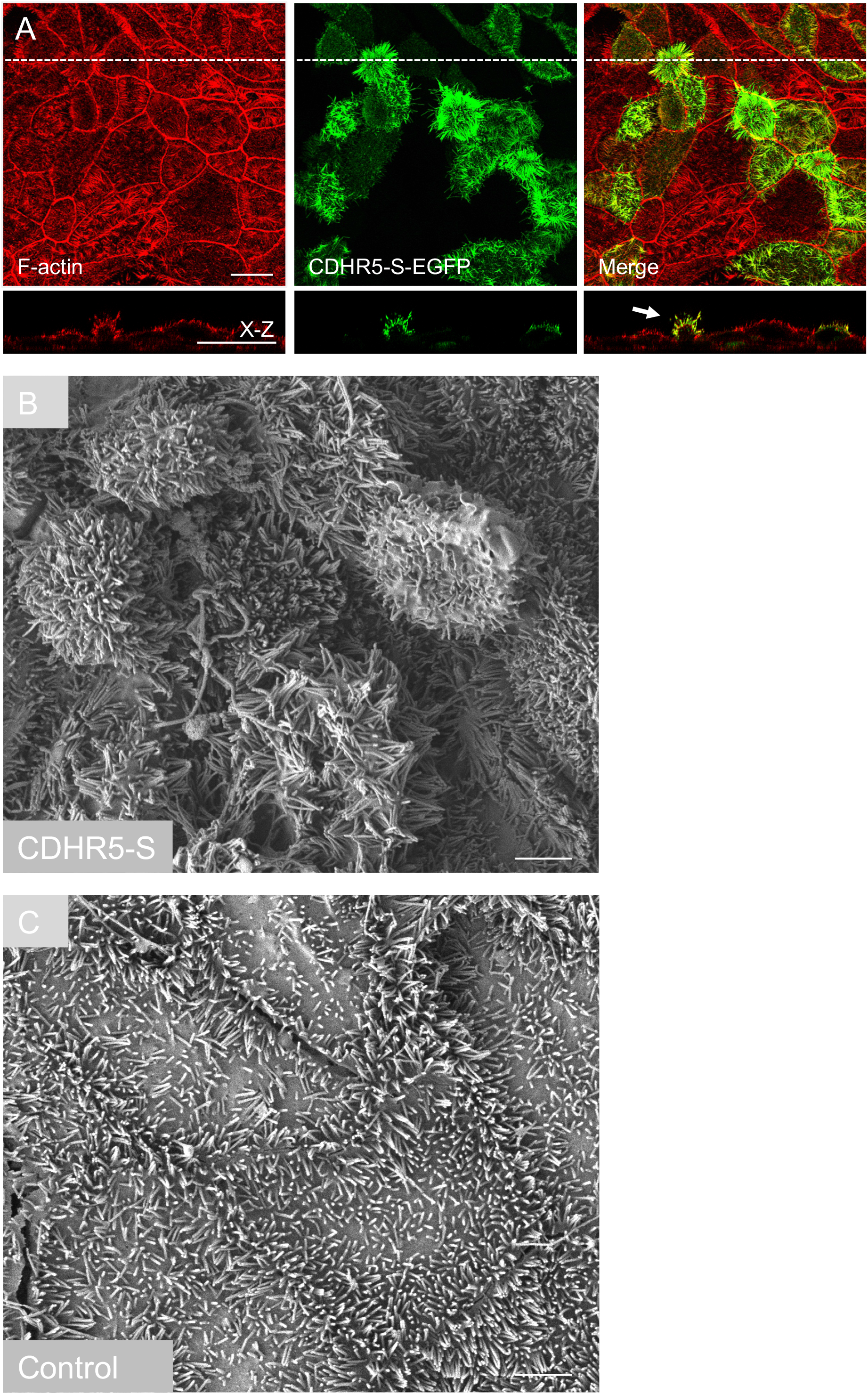
Overexpression of CDHR5-S results in changes in the surface features of LLC-PK1-CL4 monolayers. (A) Confocal images of 4-day polarized LLC-PK1-CL4 cells stably expressing EGFP-tagged CDHR5-S (green) and stained for F-actin (red). Dashed line indicates the position where the x-z section was taken; Different channels for the x-z section are shown below each *en face* image channel. Arrow points to a cell overexpressing CDHR5-S-EGFP that has a herniated apical membrane, with long microvilli splaying outwards from the herniation. Scale bars, 10 μm. (B+C) Scanning electron microscopy images of the surface features of a 4-day polarized LLC-PK1-CL4 cell monolayer either stably overexpressing CDHR5-S-EGFP or an empty vector control. Scale bars, 5 μm.

**Figure S5.**
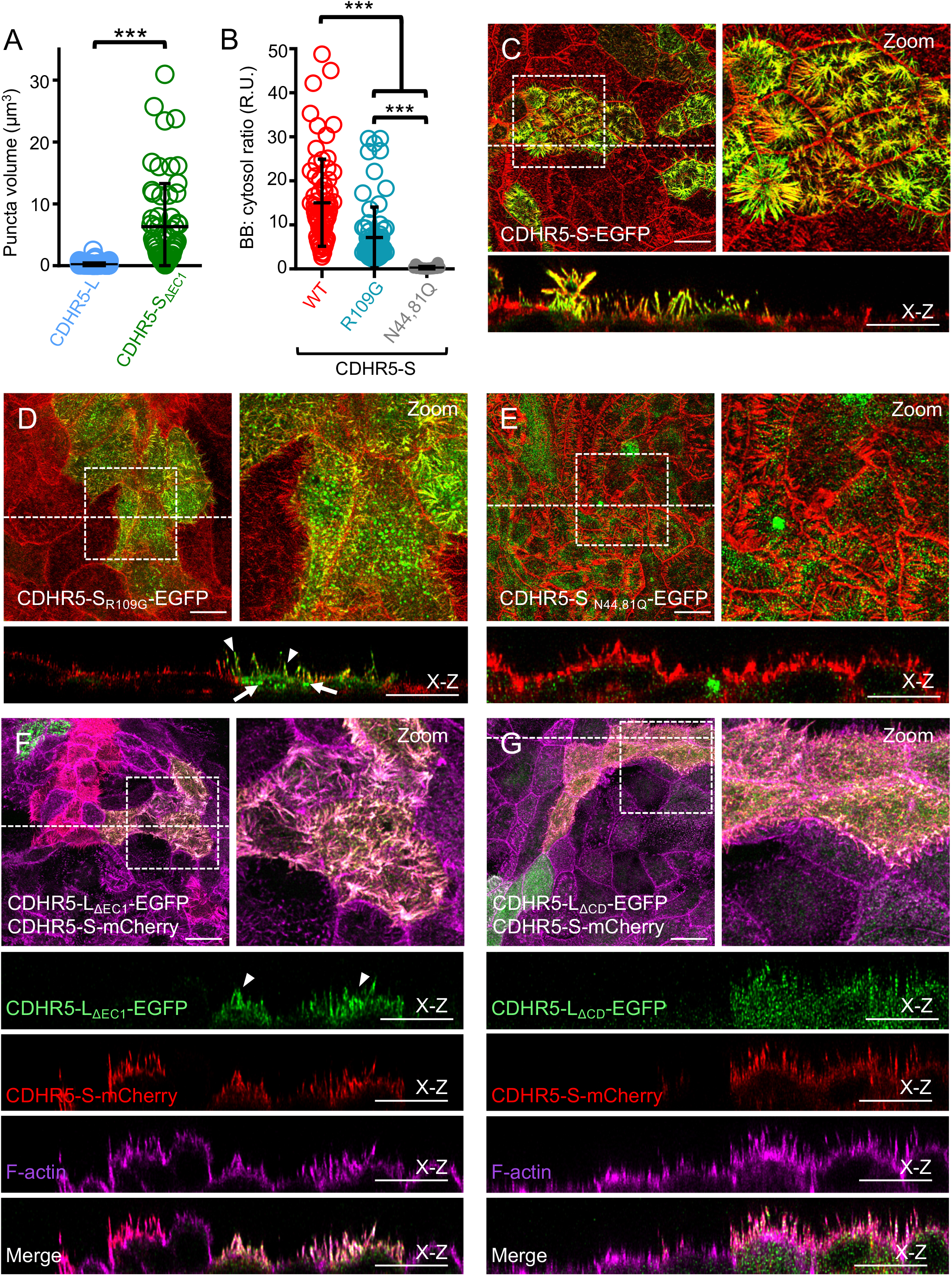
Characterization of the apical targeting properties of the splice isoforms of CDHR5 in kidney epithelial cells. (A) Scatterplot quantification of the volume of EGFP-positive puncta between LLC-PK1-CL4 cells overexpressing EGFP-tagged CDHR5-L and CDHR5-S_ΔEC1_. Bars indicate mean and SD. ***p < 0.0001, two-tailed t test. Deletion of EC1 from CDHR5-S results in formation of large intracellular EGFP-positive puncta. (B) Scatterplot quantification of the BB:cytosol ratios of EGFP signal from cells expressing wild-type CDHR5-S, the CDHR5-S_R109G_ adhesion-deficient mutant, and the CDHR5-S_N44,81Q_ glycosylation mutant. Bars indicate mean and SD. R.U = relative units. ***p < 0.0001, two-tailed t test. (C-E) Confocal images of 4-day polarized LLC-PK1-CL4 cells stably expressing EGFP-tagged constructs tested (green) stained for F-actin (red). Boxed regions denote area in zoomed image panels. Dashed lines indicate the positions where the x-z sections were taken; x-z sections are shown below each *en face* image. Arrowheads denotes localization to BB microvilli, while arrows point to signal accumulation found in subapical and cytoplasmic puncta. Scale bars, 10 μm. Both the adhesion interface and N-glycosylation may play a role in targeting CDHR5-S to apical microvilli. (F-G) Confocal images of 4-day polarized LLC-PK1-CL4 cells stably expressing both EGFP-tagged CDHR5-L variant constructs (green) and mCherry-tagged CDHR5-S (red) and stained for F-actin (magenta). Boxed regions denote area in zoomed image panels. Dashed lines indicate the positions where the x-z sections were taken; Individual channels and a merge image for the x-z section are shown below the *en face* image. Arrowheads in the CDHR5-L_ΔEC1_-EGFP x-z section green channel point to BB enrichment of signal. Scale bars, 10 μm. Co-expression of CDHR5-S promotes apical targeting of CDHR5-L_ΔEC1_, but only partial BB targeting of CDHR5-L_ΔCD._

**Figure S6.**
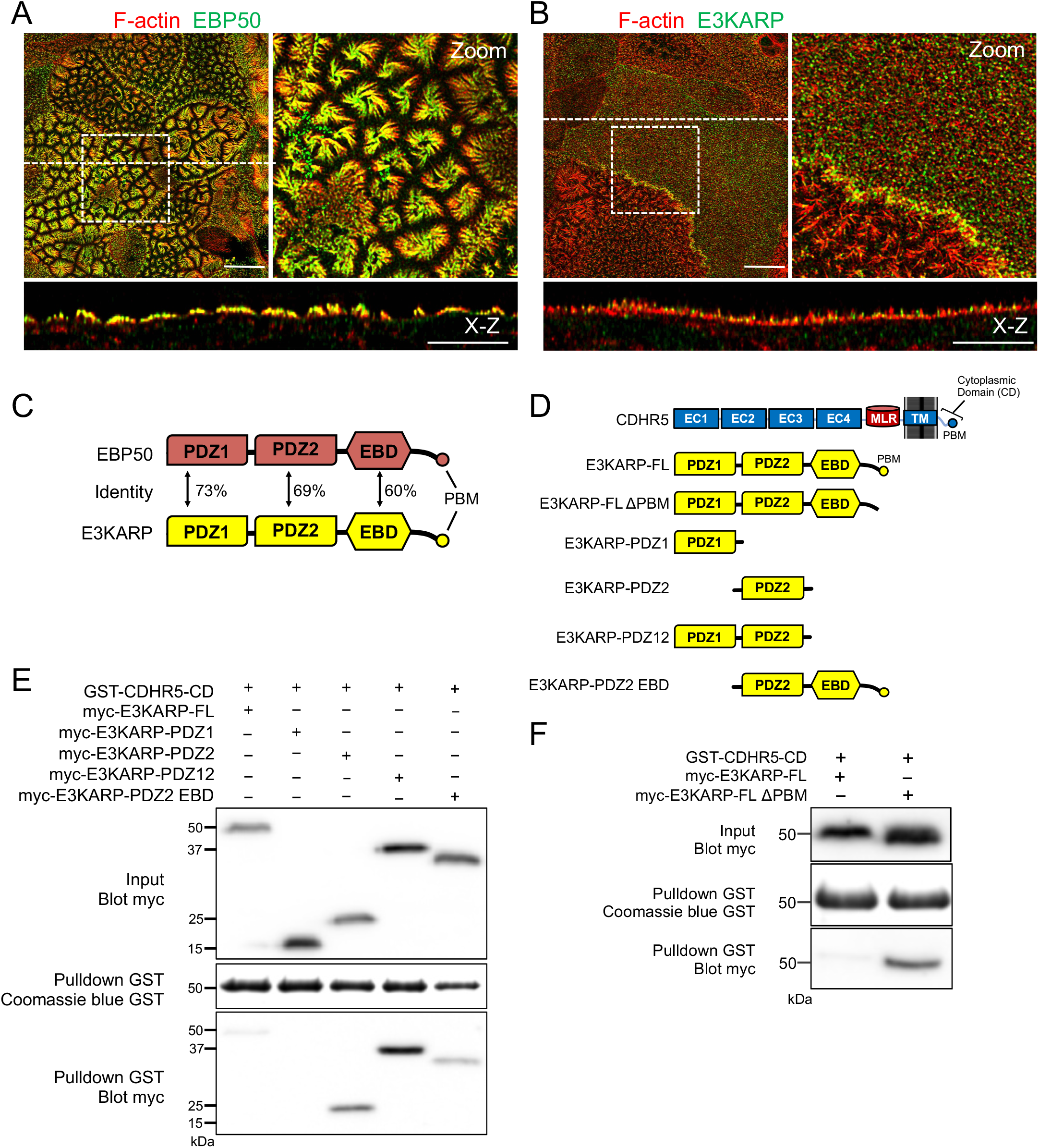
Validation of the interaction between CDHR5 and the Ezrin-associated scaffolds E3KARP and EBP50. (A-B) Confocal images of 12-day polarized CACO-2_BBE_ cells stained for F-actin (red) and either endogenous EBP50 or endogenous E3KARP (green). Boxed regions denote area in zoomed image panels. Dashed lines indicate the positions where x-z sections were taken; x-z sections are shown below each *en face* image. Scale bars, 10μm. EBP50 exhibits robust targeting to BB microvilli. (C) Domain diagram showing the sequence identity between EBP50 and E3KARP. (D) Diagram of E3KARP constructs used to map interaction with CDHR5. (E-F) Mapping the interaction between E3KARP and the cytoplasmic tail of CDHR5. Beads coated with bacterially-expressed GST-CDHR5 CD served as bait, while COS7 cell lysates expressing myc-tagged E3KARP constructs served as pulldown material containing the prey. CD= cytoplasmic domain, FL= Full-length, PDZ12=fragment comprised of PDZ1 and PDZ2, EBD=Ezrin binding domain. The cytoplasmic tail of CDHR5 interacts with the open, active version of E3KARP through PDZ2.

**Figure S7.**
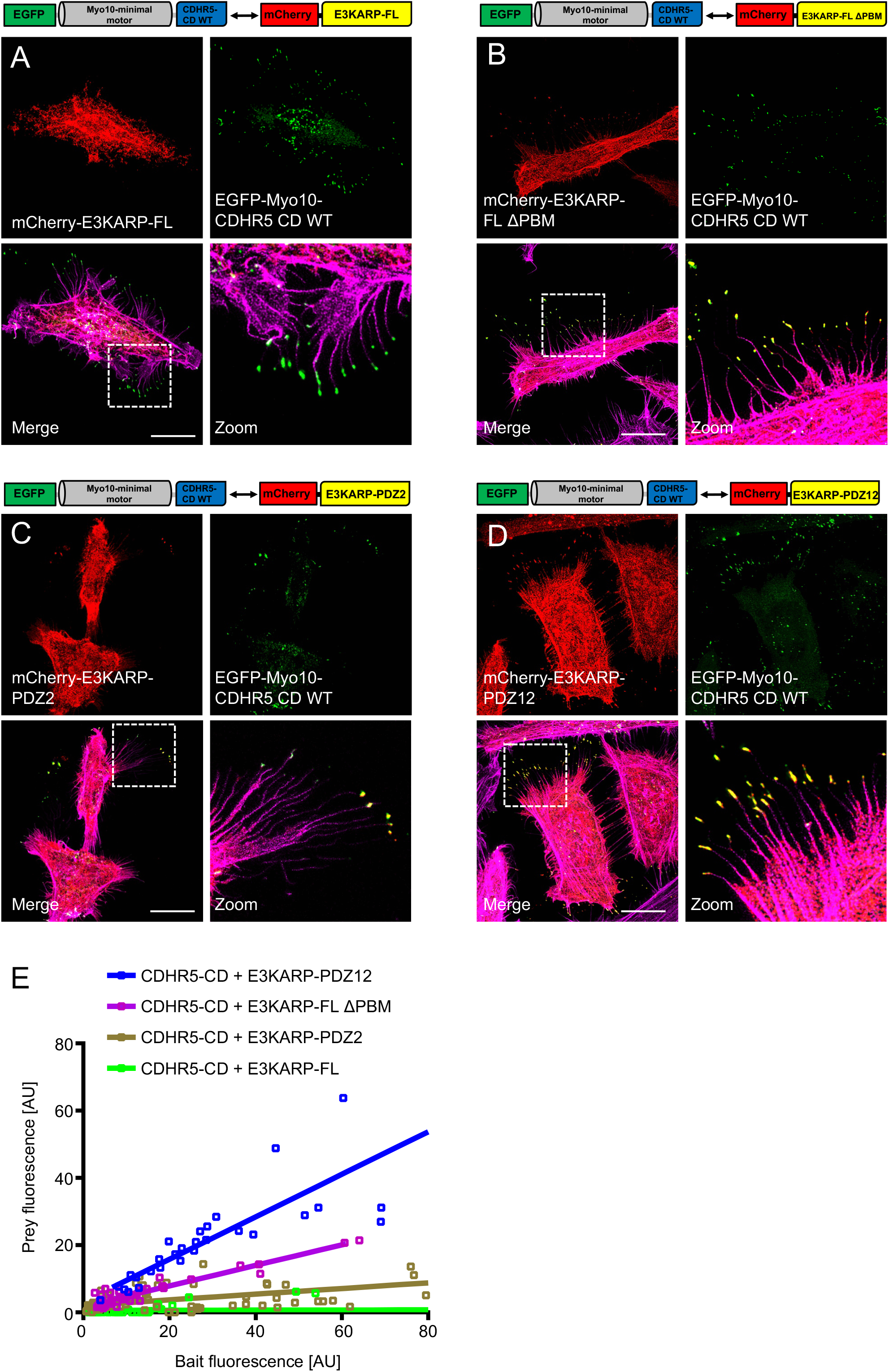
E3KARP can interact with the cytoplasmic tail of CDHR5 in a cellular context. (A-E) Confocal images of nanotrap pulldowns experiments between the cytoplasmic tail of CDHR5 and E3KARP constructs. See text for nanotrap experimental details. GFP-tagged Myo10-HMM-CDHR5 CD constructs served as bait, mCherry-tagged E3KARP constructs served as prey. Boxed regions in merge panels denote area in zoomed image panels. Scale bars,10 μm. E3KARP can interact with the cytoplasmic tail of CDHR5 in a cellular context. (F) Quantification of the correlation between bait (x-axis) and prey (y-axis) fluorescence at individual filopodia tips. Lines represent best linear fit. CD= cytoplasmic domain, FL= Full-length, PDZ12=fragment comprised of PDZ1 and PDZ2.

**Figure S8.**
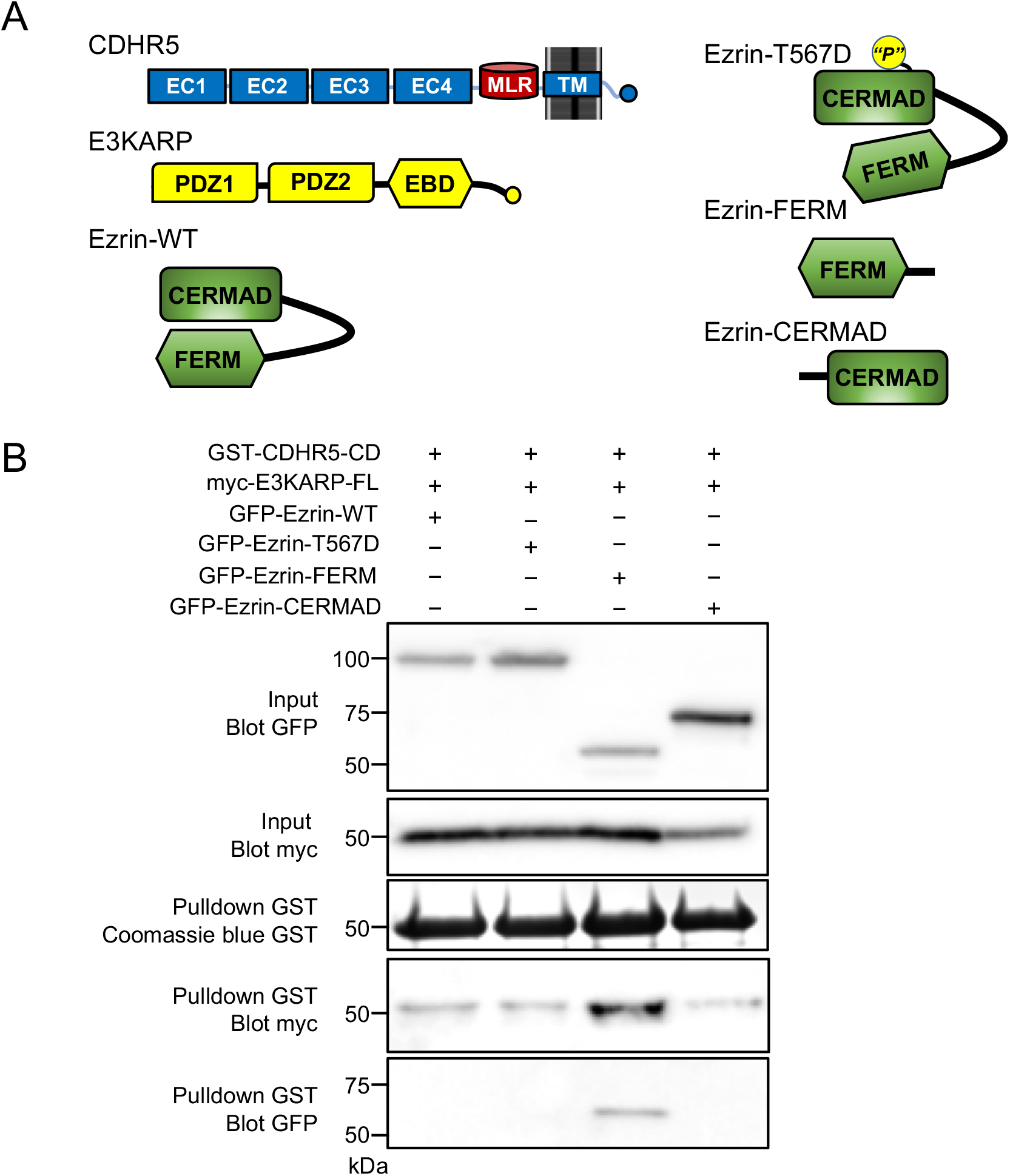
Ternary Complex Formation between CDHR5, E3KARP, and Ezrin. (A) Cartoon schematic of the different Ezrin constructs used in pulldown assays with E3KARP and the cytoplasmic tail of CDHR5. (B) Ternary complex formation between CDHR5, E3KARP and Ezrin. Beads coated with bacterially-expressed GST-CDHR5 CD served as bait, while COS7 cell lysates expressing myc-tagged E3KARP and various EGFP-tagged Ezrin constructs served as pulldown material containing the prey. A fragment of Ezrin can activate E3KARP to allow it to interact with the cytoplasmic tail of CDHR5.

**Figure S9.**
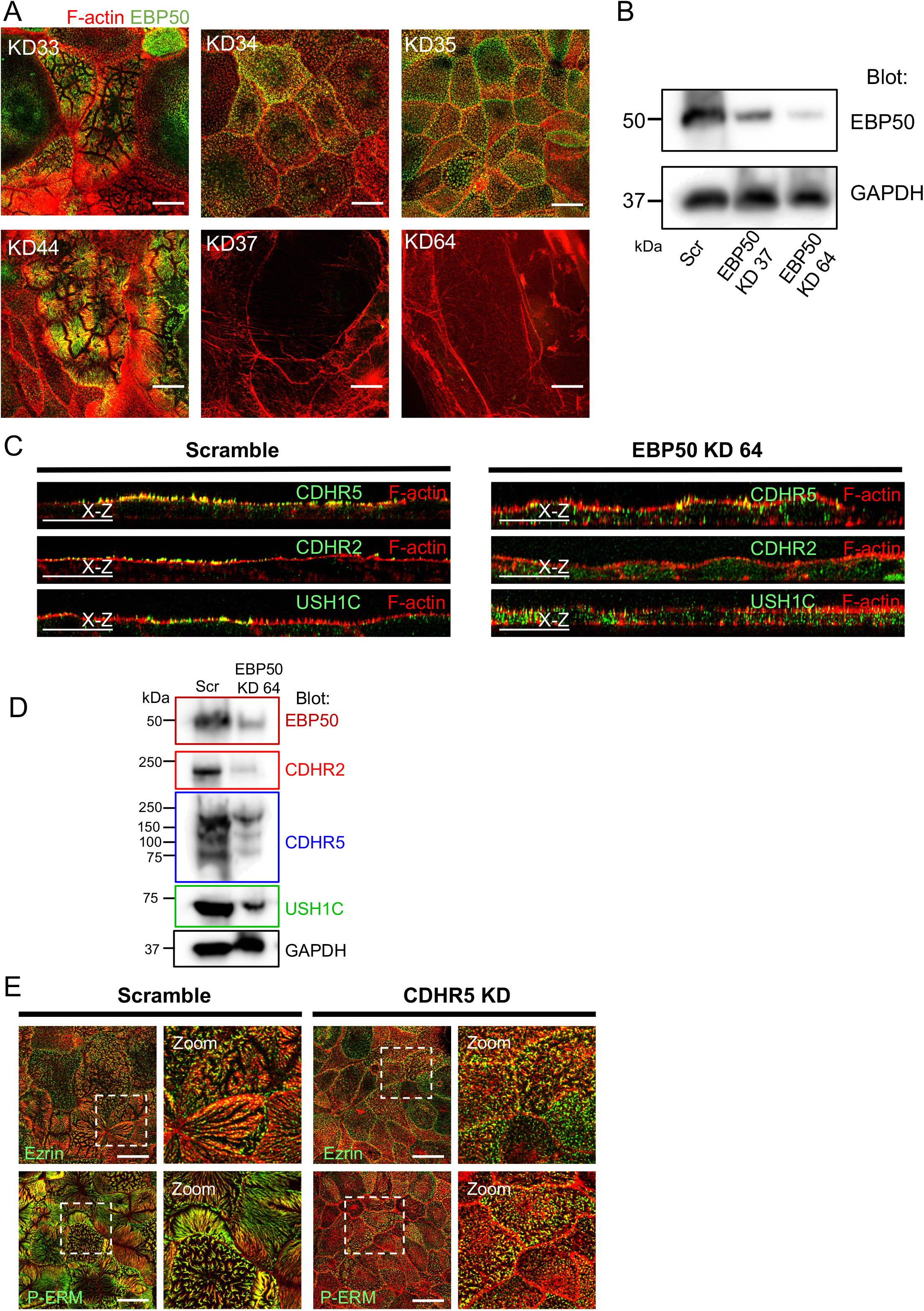
Loss of EBP50 results in lower levels of apical IMAC. (A) Confocal images of 12-day polarized CACO-2_BBE_ cells stably expressing either a scramble shRNA construct or one of six independent shRNAs targeting EBP50, stained for F-actin (red) and EBP50 (green). Scale bars, 15 μm. (B) Immunoblot analysis of endogenous EBP50 levels in lysates from 21-day polarized scramble shRNA control and two independent shRNA EBP50 KD stable CACO-2_BBE_ lines. GAPDH served as a loading control. Stable cell lines were derived three independent times for immunoblot analysis. (C) Example x-z sections taken from confocal images of 12-day polarized scramble shRNA control or shRNA EBP50 KD 64 stable CACO-2_BBE_ cell lines. Monolayers were stained for F-actin (red) and CDHR5, CDHR2, or USH1C (green) as labelled. (D) Immunoblot analysis of endogenous EBP50, CDHR5, CDHR2 and USH1C levels in lysates from 21-day polarized scramble shRNA control and shRNA EBP50 KD64 stable CACO-2_BBE_ lines. GAPDH served as a loading control. (E) Confocal images of 12-day polarized CACO-2_BBE_ cells stably expressing either a scramble shRNA construct or an shRNA targeting CDHR5, stained for F-actin (red) and either Ezrin or P-ERM (green). Boxed regions denote the area in zoomed image panels. Scale bars, 10 μm.

